# Aging restricts colorectal tumor growth by epigenetically silencing developmental gene programs

**DOI:** 10.64898/2026.06.12.731922

**Authors:** Yang Liu, Venkataramana Thiriveedi, Saratchandra Singh Khumukcham, Babak Mirminachi, Reuben R. Cano, Oladimeji Aladelokun, Sahil Choudri, Vyom Patel, Shahyan R. Khan, Shabna Mottemmal, Nicholas O. Markham, Sajid A. Khan, Caroline H. Johnson, Sara A. Grimm, Jatin Roper, Paul A. Wade

## Abstract

The incidence of early-onset colorectal cancer (CRC) has risen sharply in recent decades^1^, yet the biological basis underlying the distinct behavior of tumors arising in young versus aged tissues remains poorly understood. Here we show that aging reprograms the epigenetic landscape of the colon, restricting colon tumor growth through stable silencing of developmental and fetal gene programs. We find that colon tumors arising in aged mice are intrinsically less proliferative than those arising in young animals. Multi-omic profiling of normal colon and colon tumors reveals that aging drives DNA hypermethylation, loss of Polycomb-associated chromatin states, and reduced chromatin accessibility at a defined set of developmental genes that are bivalent (marked by both H3K27me3 and H3K4 methylation), transcriptionally active in colon tumors from young animals and repressed in both tumors and normal tissue from old animals. Among the genes most strongly repressed in old animals is *Tacstd2* (Trop2), a regulator of fetal intestinal programs and epithelial stemness. Pharmacologic inhibition of DNA methylation reactivates the aging-silenced gene network in organoids from old animals, whereas genetic disruption of *Tacstd2* suppresses growth and developmental transcriptional programs in young tumor organoids. *TACSTD2*, fetal gene signatures, and the aging-associated bivalent gene program are likewise repressed in late-onset vs. early-onset human colorectal cancers. Collectively, these findings identify age-associated epigenetic silencing of developmental gene programs as a causal mechanism that constrains colorectal tumor growth and provide a mechanistic framework for understanding the distinct biology of early-onset colorectal cancer.

## MAIN

Colorectal cancer (CRC) is among the most common and lethal malignancies worldwide^2^. Early-onset CRC, diagnosed before age 50, is increasing in incidence and is associated with distinct molecular features and poorer prognosis compared to late-onset disease, suggesting that age shapes tumor biology beyond accumulating somatic mutations^1,3,4^. Aging is accompanied by progressive remodeling of the epigenome, including widespread gain of DNA methylation at Polycomb-regulated loci in the normal colonic epithelium, a phenomenon that has been proposed to prime cells for malignant transformation, but whose functional consequences for tumor growth remain unclear^5–8^. Epigenetic plasticity, particularly the reactivation of fetal and developmental gene programs, has emerged as a key driver of intestinal tumorigenesis^9–11^. However, it is unknown whether a mechanistic link exists between age-associated epigenetic changes and reactivation of developmental gene programs during tumor initiation.

### Aging intrinsically limits colorectal tumor growth

To determine how aging influences colorectal tumorigenesis, we induced colon-specific *Apc* deletion by colonoscopy-guided delivery of 4-hydroxytamoxifen in 2–3-month-old (young) and 18–22-month-old (old) *Apc^fl/fl^;Villin^CreERT2^* mice, resulting in the formation of precancerous adenomas (Fig. 1A). Tumor burden, assessed by endoscopic measurements, gross tumor area at necropsy, and tumor weight was significantly reduced in aged mice compared with young controls (Fig. 1B–E). To determine whether this phenotype is tumor cell-intrinsic, EpCAM⁺ tumor epithelial cells were isolated by fluorescence-activated cell sorting and cultured as organoids. Organoids derived from old tumors exhibited reduced proliferation, smaller size, and reduced clonogenic efficiency relative to organoids derived from young tumors (Fig. 1F-I). These differences persisted following passage (Fig. 1J–K). Collectively, these findings indicate that aging attenuates tumor growth via an epithelial-intrinsic mechanism, consistent with epigenetic regulation.

**Figure 1.**
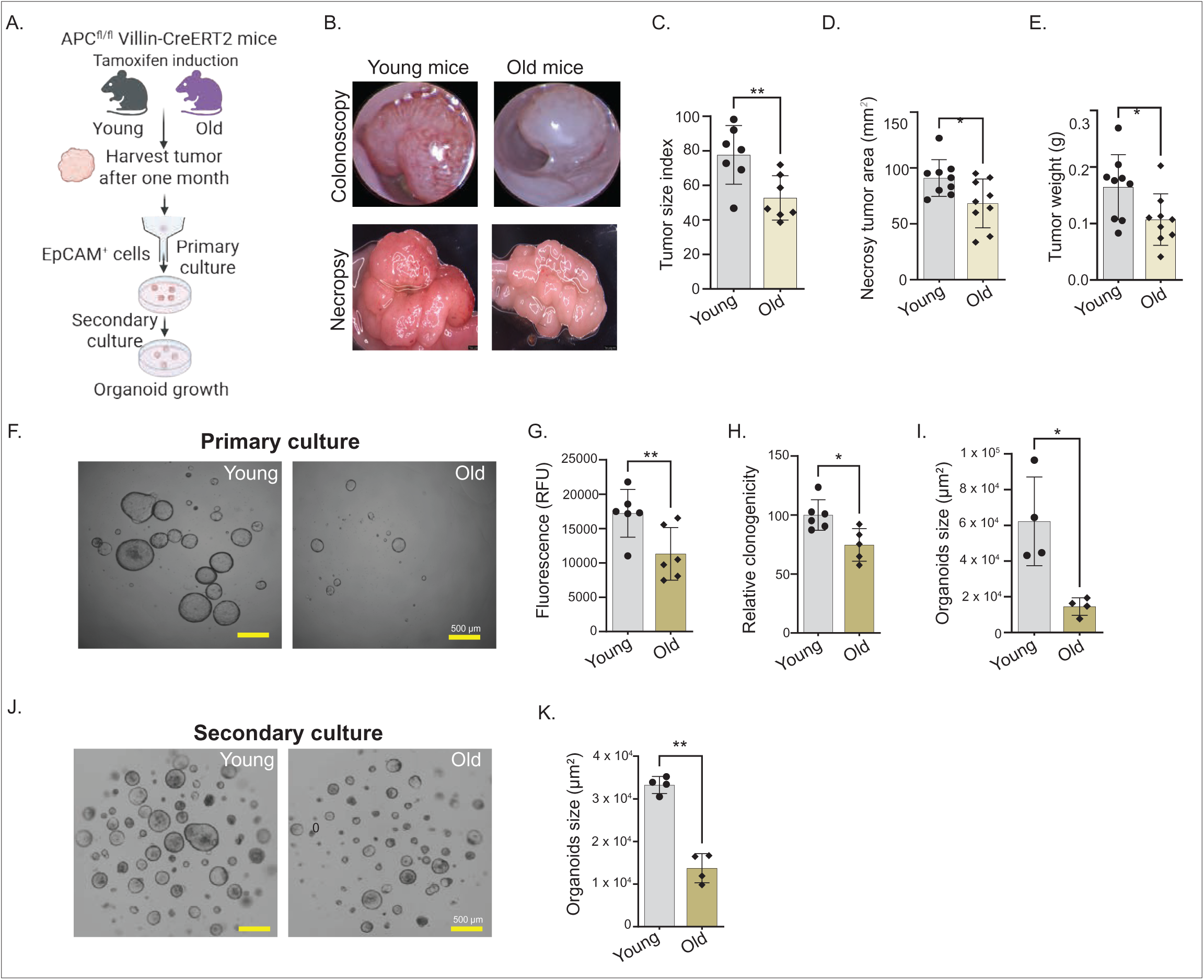
Aging alleviates colorectal tumorigenesis in an autochthonous genetically engineered mouse model. (A) Schematic of the experimental design. *Apc^fl/fl^ Villin^CreERT2^* mice (young and old) were administered tamoxifen via colonoscopy-guided delivery to induce colonic tumorigenesis. Tumors were harvested after one month for analysis. EpCAM⁺ tumor epithelial cells were isolated and cultured to evaluate primary and secondary organoid formation. (B) Representative colonoscopy and necropsy images reveal reduced tumor burden in old mice. (C–E) Quantification of tumor burden by colonoscopy-based tumor size index (C), gross tumor area at necropsy (D), and tumor weight (E); n = 7–9 mice per group. (F) Representative brightfield images of primary tumor organoids derived from EpCAM⁺ cells isolated from young and old mice. (G–I) Quantification of primary organoid-forming efficiency based on fluorescence intensity (G), clonogenic potential (H), and average organoid size (I). (J) Representative brightfield images of secondary organoids derived from passaged primary tumor cultures. (K) Quantification of secondary organoid size. Data are presented as mean ± s.e.m. Statistical significance was determined using an unpaired two-tailed t-test; P < 0.05, P < 0.01. Scale bars, 500 µm.

### Age-Associated DNA Hypermethylation Precedes Oncogenic Transformation at Polycomb-Regulated Loci

To investigate the molecular basis of these age-associated differences, we performed array-based DNA methylation profiling of epithelial crypts, and tumors from tumor-bearing young and aged mice. Consistent with prior reports describing age-associated accumulation of DNA methylation at regulatory regions in the colon^5,6,8^, we identified 7,261 CpG sites with increased methylation in colon tumors from aged versus young mice (Figure 2A) and a similar number (7821) of hypermethylated CpG sites, with substantial overlap, in colon crypts from aged versus young animals (Figure 2B, Extended Data Figure 1A). This overlap between age-associated hypermethylation in normal and tumor tissue suggests that these changes in epigenetic status precede oncogenic transformation from colonic stem cells and do not arise through tumor-specific selection. As previously reported^5^, age-dependent hypermethylation events are enriched for DNA motifs characteristic of Polycomb Repressive Complex binding (Extended Data Figure 1B-C) in both tumor and normal tissue, implicating chromatin-based regulatory mechanisms. We next selected a locus exemplifying this pattern, *Tacstd2* for more detailed scrutiny. Methylation levels of CpG dinucleotides within the *Tacstd2* promoter region illustrate elevated DNA methylation in both tumors and normal crypts from aged compared with young animals, illustrating that this developmental regulatory locus undergoes age-associated hypermethylation (Figure 2C).

**Figure 2.**
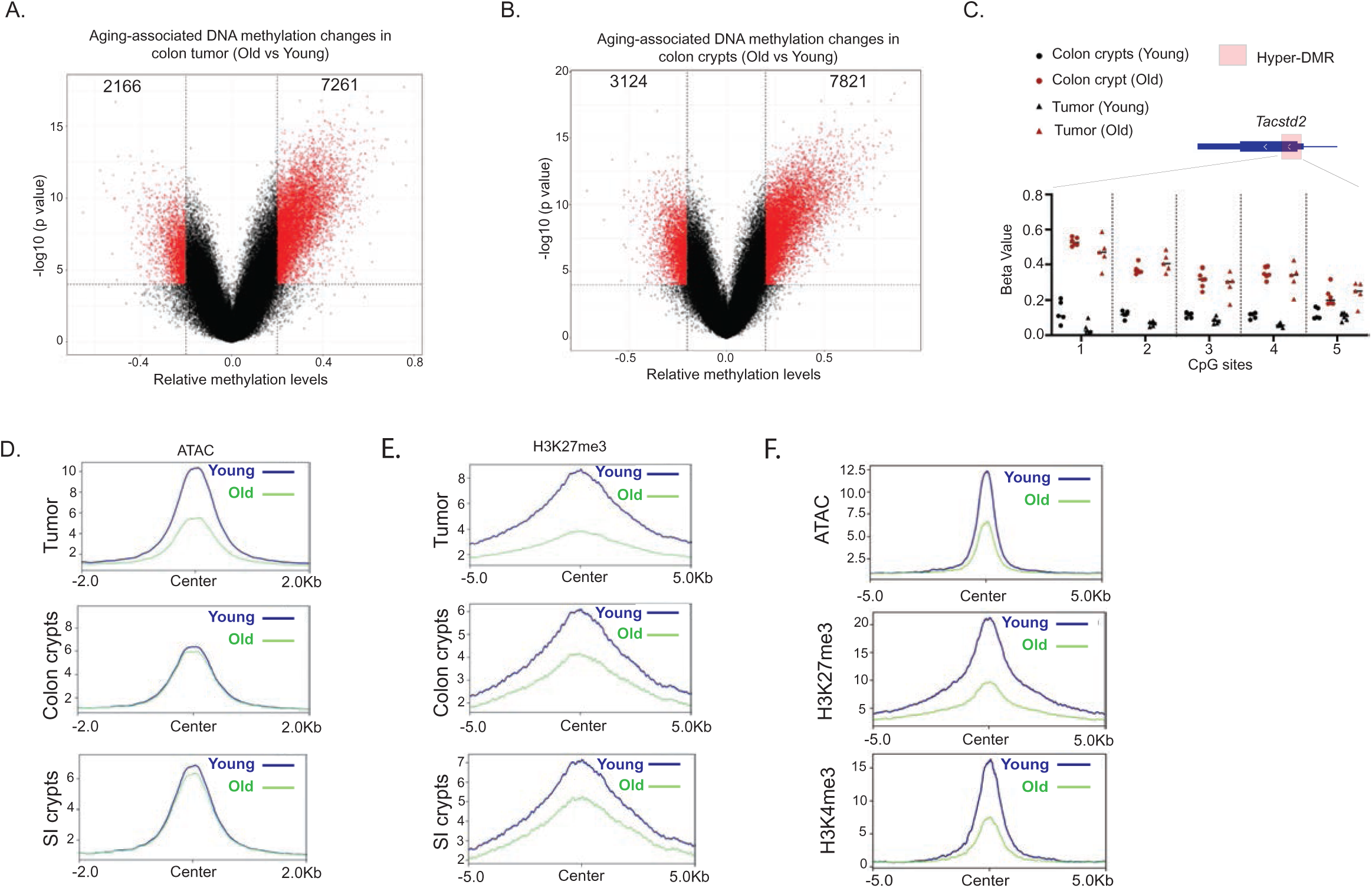
Age-dependent alterations in epigenetic marks in normal colon and tumor tissue. (A) The volcano plot depicts differentially methylated positions (DMPs) in tumors during aging. (B) The volcano plot depicts differentially methylated positions (DMPs) in normal colonic crypts during aging. (C) DNA methylation status at individual CpG residues at the *Tacstd2* locus in normal and tumor tissue from old and young individuals are depicted as Beta value. Approximate genomic locations of these CpG sites are indicated in the cartoon above the data. (D) The metagene plots depict the levels of chromatin accessibility as assessed by ATAC-seq for the set of loci with decreased accessibility in tumors from old animals. Data depict the aggregate value for tumor tissue, matched normal colon crypts, and matched normal small intestine. (E) The metagene plots indicate the levels of H3K27me3 modification at the set of loci displaying decreased accessibility by ATAC in tumors from old animals. Data are shown for tumor tissue, matched normal colon crypts and matched normal small intestine. (F) The metagene plots depict the aggregate levels of chromatin accessibility and the indicated histone modifications at loci exhibiting decreased accessibility in old tumors that are marked by H3K4me3.

### Aging Drives Widespread Changes in Chromatin Accessibility and Histone Modifications in Colorectal Tumors

Alterations in chromatin accessibility have also been linked to aging^12–15^. Therefore, we next performed the Assay for Transposase Accessible Chromatin using sequencing (ATAC-seq^16^) to assess whether chromatin accessibility changes with age. We identified numerous loci (1677) exhibiting decreased chromatin accessibility in tumors from old compared to young animals, whereas accessibility at these loci remained unchanged with aging in matched normal small intestine and colon (Figure 2D, Extended Data Figure 1D). Notably, a significant number of age-dependent differentially accessible regions overlapped with loci that gained DNA methylation in old versus young tumors, linking age-associated DNA hypermethylation to reduced accessibility during tumor initiation (Extended Data Figure 1E).

Age-dependent DNA methylation in normal colon has been previously linked to chromatin state^8,17^. To assess the impact of aging on promoter chromatin state in our mouse model of colorectal cancer initiation, we performed CUT&TAG (Cleavage Under Targets and Tagmentation^18^) for trimethylation of lysine 4 of histone H3 (H3K4me3, which marks regions competent for transcription) and trimethylation of lysine 27 of histone H3 (H3K27me3, which marks transcriptionally repressed loci) in tumor and matched normal tissue from young and aged mice. Overall, we observed widespread loss of H3K27me3 enrichment with age in both tumor and normal intestinal/colon crypts and EpCAM+ cells (Figure 2E, Extended Data Figure 2A-B). In addition, age-dependent DNA hypermethylation in tumors was significantly negatively correlated with H3K27me3 levels (Extended Data Figure 2C).

### Integrative Epigenomic Analysis Identifies a Developmentally Regulated Gene Set Subject to Age-Dependent Silencing

To identify promoters affected by aging, we intersected H3K4me3-marked regions in young tumors with loci exhibiting decreased chromatin accessibility in tumors from aged mice, identifying approximately 1000 regions. Of these, 85% also carried H3K27me3 in young tumors and normal tissue, a significant enrichment compared with background regions (Fisher’s exact test, p<0.0001). These regions were predominantly marked by H3K27me3 in young tumors, consistent with a transition to bivalency after tumor initiation (Figure 2F, Extended Data Figure 3A-B). This chromatin state pattern was also observed small intestinal crypts and in sorted EpCAM+ tumor cells (Extended Data Figure 3C-E).

We next performed RNA sequencing on tumor and matched normal intestinal/colonic crypts from young and aged mice. Approximately 150 genes were expressed at higher levels in tumors from young animals and a similar number were expressed at higher levels in tumors from aged animals (Figure 3A), with comparable age-associated differences observed in matched normal crypts (Extended Data Figure 4A). Integrating transcriptomic, DNA methylation, chromatin accessibility, and histone modification data, we identified a core set of 46 genes that were (i) strongly differentially expressed between aged and young tumors but uniformly repressed in normal tissue (Figure 3B), (ii) bivalent in young tumors (Figure 3C), (iii) exhibited decreased chromatin accessibility in old versus young tumors (Figure 3C), and (iv) gained DNA methylation during aging (Figure 3D). The epigenetic features of the *Tacstd2* locus in tumors and matched normal crypts exemplify the features of this gene set (Figure 2C, Figure 3E).

**Figure 3.**
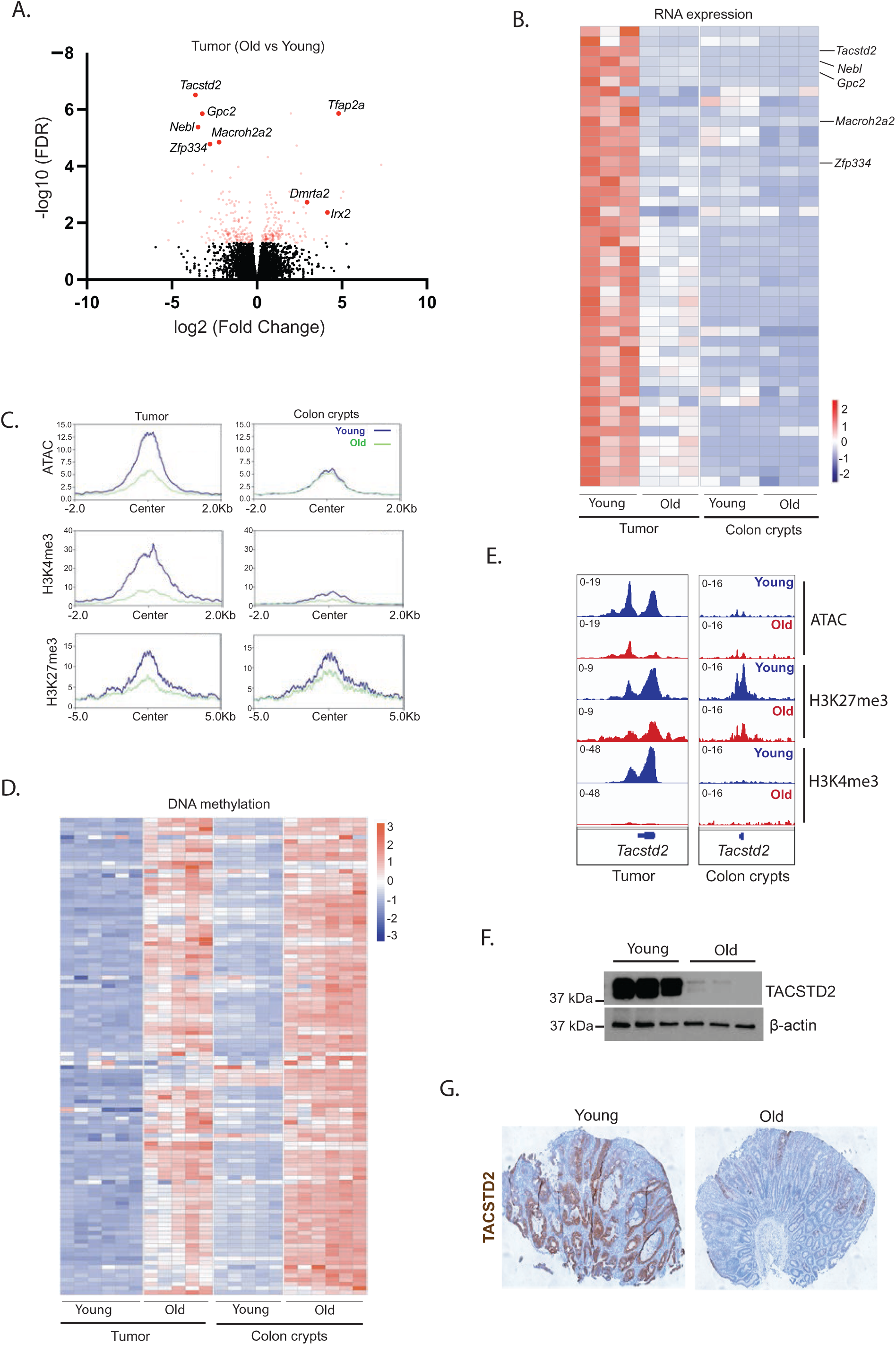
Integration of multiple high-content data streams identifies a genes distinguishing tumors in old animals versus young animals. (A) The volcano plot depicts transcripts differentially expressed in old versus young tumors. Genes of interest are highlighted in the plot. (B) The heat map depicts expression of the 46 gene set described in the text in tumors and matched normal crypts from old and young animals. (C) The metagene plots show aggregate levels of chromatin accessibility and the indicated histone modification at chromatin-accessible regions associated with the 46-gene set in tumors and matched normal crypts from young and old animals. (D) Heat map showing DNA methylation levels at chromatin-accessible regions associated with the 46-gene set in tumors and matched normal crypts from young and old animals. (E) IGV tracks showing indicated histone modifications enrichment and chromatin accessibility at *Tacstd2* in young and old tumors and matched normal crypts. (F) Immunoblot of TACSTD2 in *Apc*-null colon organoids from young (8–12 weeks) and old (18–24 months) mice. β-Actin, loading control. *n* = 3 independent organoid lines per group. (G) Representative TACSTD2 immunohistochemistry of *Apc* null tumors from young and old mice. *n* = 6 mice per group. Scale bars, 100 µm.

Most of these genes are downstream targets of *Wnt* signaling, an essential pathway in intestinal stem cell self-renewal and colorectal cancer initiation^19^ (Extended Data Table 1). These genes are also enriched in developmental processes, patterning, ECM remodeling and transcriptional regulation (Extended Data Table 2). Further, many members are expressed in the fetal colon^20^ (Extended Data Table 3). Together these findings identify a developmentally regulated gene program that is epigenetically poised in young tumors but becomes constrained by age-associated chromatin remodeling and DNA hypermethylation.

Overall, these data support a model in which genes expressed during fetal development become repressed in normal intestinal epithelium. Following oncogenic insult, these genes transition to a bivalent, transcriptionally permissive state in tumors from young animals, allowing partial reactivation of fetal/developmental programs. In aged animals, these same loci show reduced chromatin accessibility, diminished Polycomb-associated regulation, and acquisition of DNA methylation, consistent with a shift toward more stable transcriptional silencing.

To evaluate the role of the aging host microenvironment in these processes, we implanted AKPST CRC organoids^21^ (*Apc*Δ/Δ;*Kras*^G12D/+^;*Trp53*Δ/Δ null;*Smad4*Δ/Δ) into young and old wild-type mice (Extended Data Figure 5A). Unlike *Apc*-null tumors, AKPST tumors showed minimal aging-associated transcriptomic and epigenetic changes, including at the *Tacstd2* locus (Extended Data Figure 5B–D), suggesting a limited contribution of the aging microenvironment to tumor epigenetic alterations.

### *Tacstd2* is a Hallmark of the Age-Dependent Epigenetic Program

Among the most differentially expressed genes in this group was *Tacstd2* (Tumor-Associated Calcium Signal Transducer 2, encoding the Trop2 protein), which is overexpressed in multiple human cancers^22^, is epigenetically regulated in human colorectal tumors^23,24^, and marks a fetal intestinal gene expression program reactivated following *Apc* loss^9^. Quantitative PCR analysis of sorted EpCAM^+^ *Apc*-null tumor cells (Extended Data Figure 6A), western blot of tumor organoids (Figure 3F), and immunohistochemistry of tumor sections (Figure 3G, Extended Data Figure 6B) demonstrated significant repression of *Tacstd2*/Trop2 in tumors from aged versus young mice.

### DNA Methylation Contributes to Epigenetic Silencing of Stemness-Associated Gene Networks in Aged Tumors

To assess the functional significance of age-dependent epigenetic changes, we used organoids derived from *Apc*-null tumors. EpCAM+ tumor cells isolated from aged mice were embedded in Matrigel and treated with a Dnmt1 inhibitor to reduce global DNA methylation, or vehicle control (Figure 4A), followed by RNA sequencing after four days. Among the genes upregulated by Dnmt1 inhibition, 17 members of our 46 gene set identified by epigenomic analysis were significantly induced, including *Tacstd2* (Figure 4B–D), demonstrating that age-dependent epigenetic silencing suppresses this gene set in tumors from aged animals. Pathway analysis of genes activated by Dnmt1 inhibition revealed a dominant immune and inflammatory transcriptional response, consistent with derepression of endogenous retroviruses and the viral mimicry mechanism (Extended Data Table 4). In addition, enrichment of epithelial-to-mesenchymal transition and Reactome terms including laminin interactions and hemidesmosome assembly point to broad reactivation of epithelial identity programs. Strikingly, multiple lines of evidence indicate reactivation of *Wnt* signaling: increased expression of *Wnt* ligands (*Wnt4, Wnt7b, Wnt9a*) and co-receptors (*Gpc1, Gpc6*), induction of canonical *Wnt* transcriptional targets^25^ (*Lgr6, Runx2, Sphk1*), and upregulation of *Wnt*-regulated extracellular matrix genes required for intestinal stem cell niche integrity^26^ (*Lama3, Lamb3, Lamc2*) (Extended Data Figure 7A). These findings demonstrate that gene networks central to colon epithelial stemness are subject to age-dependent epigenetic silencing through DNA methylation.

**Figure 4.**
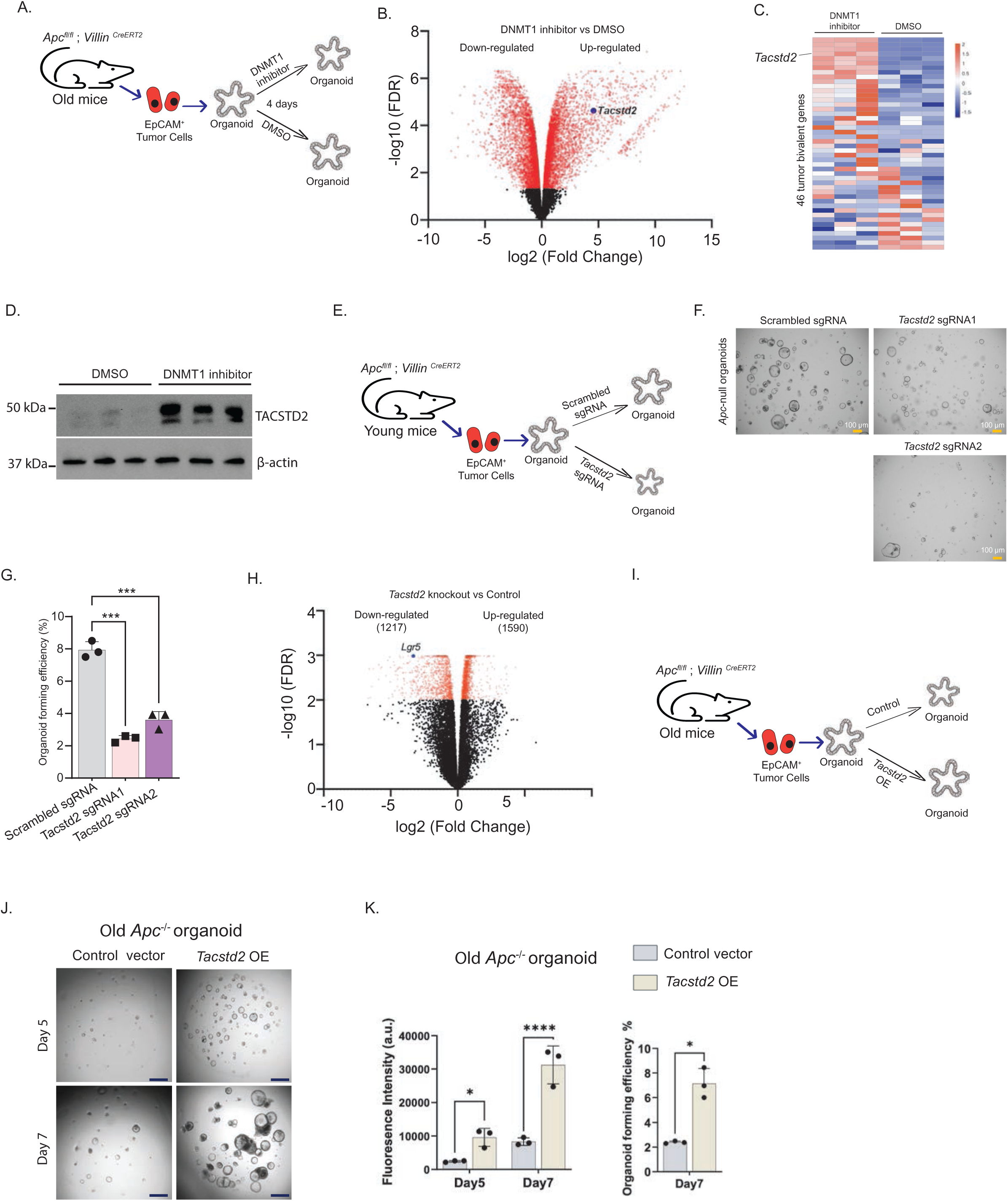
*Tacstd2* is epigenetically regulated with age. (A) Schematic diagram of the DNMT1 inhibition experiment described in the text. (B) Volcano plot showing the transcriptomic changes between DMSO and DNMT1 inhibitor treatment. *Tacstd2* is indicated in the plot. (C) Heatmap showing the expression levels of the 46 bivalent genes in the DNMT1 inhibition experiment. *Tacstd2* is indicated in the heat map. (D) Representative immunoblot of TACSTD2 in old *Apc*-mutant mouse organoids treated with the 2 µM DNMT1 inhibitor, GSK3484862, or DMSO (vehicle) for 4 days. Each lane represents an independent organoid line. β-Actin, loading control. Representative of *n* = 3 independent experiments. (E) Schematic of the *Tacstd2* genome engineering experiment utilizing tumor organoids derived from young mice. (F) Representative bright-field images of young *Apc*-null colon organoids after CRISPR–Cas9-mediated knockout of *Tacstd2* or non-targeting control (scrambled sgRNA). Scale bars, 100 µm. (G) Organoid-forming efficiency, quantified at day 4 after seeding. Data are mean ± s.d.; *n* = 3 independent biological replicates. *\*\*\*P < 0.0001;* one-way ANOVA with Tukey’s multiple comparisons test. (H) Volcano plot showing the transcriptomic changes between control and *Tacstd2* deletion organoids. *Lgr5* is indicated in the plot. (I) Schematic of the *Tacstd2* overexpression experiment utilizing tumor organoids derived from old mice. (J) Representative bright-field images of old *Apc*-null colon organoids transduced with control vector or *Tacstd2*-overexpression construct (*Tacstd2* OE), imaged at days 5 and 7 after seeding. Scale bars, 100 µm. (K) Quantification of organoid growth. Left, fluorescence intensity at days 5 and 7 (\**P* < 0.05, \*\*\**P* < 0.0001); two-tailed Student’s *t*-test; right, organoid-forming efficiency at day 7 (\**P* < 0.05), two-tailed unpaired Student’s *t*-test. Data are mean ± s.d.; *n* = 3 independent biological replicates.

### *Tacstd2* is a Functional Regulator of Colon Tumor Growth and Stemness

To determine the functional consequences of manipulating a representative member of this gene set, we genetically disrupted *Tacstd2* in organoids derived from young mice (Figure 4E). Introduction of two independent sgRNAs resulted in marked depletion of Trop2 protein (Extended Data Figure 8A-B). *Tacstd2*-deficient organoids exhibited impaired growth, reduced clonogenic capacity (Figure 4F-G) and extensive transcriptome remodeling (Figure 4H). Downregulated genes are enriched for developmental processes and transcriptional programs, including the *Wnt* signaling pathway and other biological pathways integral to stemness (Extended Data Figure 8C). Prominent amongst downregulated genes was the *Wnt* target gene *Lgr5* (Figure 4H, Extended Data Figure 8D), suggesting that sustained expression of *Tacstd2* supports a developmentally regulated gene expression network related to stemness.

Conversely, we overexpressed *Tacstd2* in tumor organoids derived from old mice, in which endogenous *Tacstd2* is epigenetically repressed (Figure 4I, Extended Data Figure 8E). Organoids overexpressing *Tacstd2* exhibited increased organoid-forming efficiency and organoid size (Figure 4J-K, Extended Data Figure 8F), indicating that restoration of age-silenced Tacstd2 enhances growth potential in aged tumor organoids. Collectively, these data identify Tacstd2 as a functional component of the age-regulated developmental program that supports colon tumor organoid growth, clonogenicity, and stemness-associated transcriptional states.

### Tumor-bivalent and fetal transcriptional programs distinguish Early-Onset from Late-Onset Colorectal Cancer

Finally, we asked whether the aging-related epigenetic remodeling of tumor bivalent and fetal/developmental transcriptional programs that we observe in a genetically engineered mouse model is relevant to human CRC. Consistent with our animal model, analysis of publicly available human CRC bulk RNA sequencing datasets^27^ revealed age-associated repression of *TACSTD2* mRNA levels (Figure 5A), the tumor bivalent gene signature identified in this work (Figure 5B), as well as a previously reported fetal intestine gene signature^28^ (Figure 5C) in human CRC. Notably, Kaplan-Meier analysis demonstrated improved survival among patients whose tumors exhibited lower *TACSTD2* expression (Figure 5D). Likewise, the *WNT*-responsive gene panel identified here was differentially expressed in human Early-Onset vs Late-Onset CRC^29^ (Figure 5E). These analyses are consistent with reactivation of a bivalent and fetal/developmental gene expression program in Early-Onset human CRC similar to the pattern observed in our mouse model.

**Figure 5.**
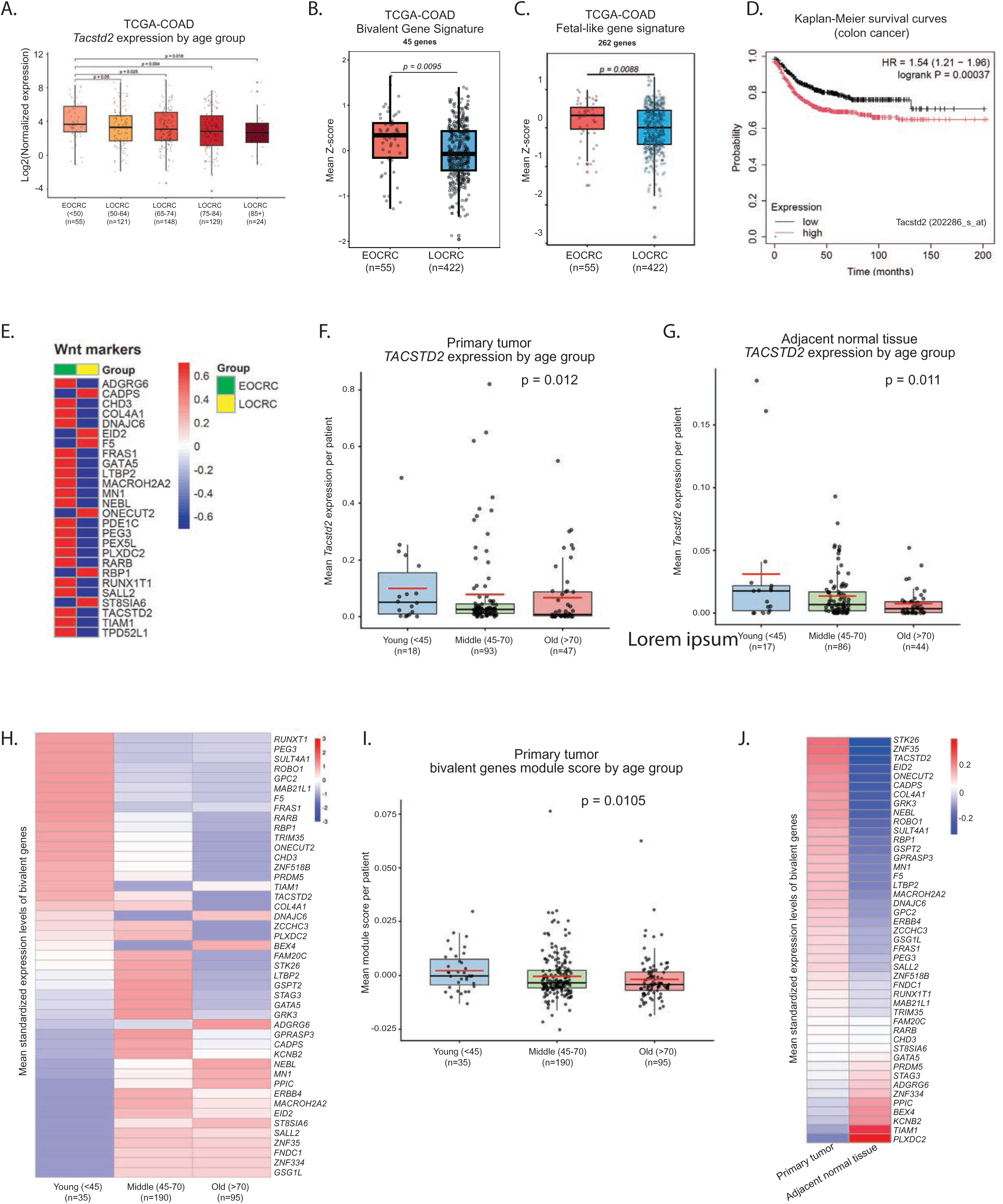
Aging-related epigenetic repression of *TACSTD2* blunts colorectal tumorigenesis. (A) The box and whisker plot depicts *TACSTD2* RNA expression in human colorectal tumors as a function of age (** represents p<0.005, using unpaired, non-parametric Wilcoxen test). Log_2_-transformed values of *TACSTD2* raw count in a total of 55 EOCRC tumors are compared with 422 LOCRC tumors after normalization to normal colon tissues in TCGA-COAD^27^ dataset. (B) Comparison of bivalent genes expression levels between EOCRC and LOCRC patients. Box plots depict the mean z-score per sample representing the bivalent signature. Differences between the bivalent signature for EOCRC and LOCRC were determined by Student’s t-test. (C) Comparison of a fetal-like gene signature^28^ (containing 262 genes) between EOCRC and LOCRC patients. Box plots depict the mean z-score per sample representing the fetal-like gene signature. Differences between the fetal-like signature for EOCRC and LOCRC were determined by Student’s t-test. (D) Kaplan-Meier survival curves for individuals with low and high *TACSTD2* mRNA expression levels. (E) Heatmap showing the expression levels of Wnt signaling markers in EOCRC and LOCRC patients. (F) Per-participant *TACSTD2* expression in primary tumor cells from Marteau et al^30^, stratified by age group (young <45, n = 18; middle 45–70, n = 93; old >70, n = 47). Each point represents one individual (mean log-normalized expression across that patient’s cells); boxes show the median and interquartile range, whiskers the 1.5× IQR, and the red crossbar the group mean. Spearman correlation, ρ ≈ −0.20, p = 0.012). (G) As in (F), for participant-matched adjacent normal tissue (young, n = 17; middle, n = 86; old, n = 44). Spearman correlation ρ ≈ −0.21, p = 0.011. (H) Heatmap of the bivalent-gene signature in primary tumor samples from Marteau et al^30^, showing per-gene mean standardized (z-scored across participants) expression collapsed by age group (young, n = 35; middle, n = 190; old, n = 95). Rows are bivalent genes (hierarchically clustered); columns are age groups shown in fixed young→old order. Red indicates higher relative expression, blue lower. (I) Per-patient bivalent-gene module score in primary tumor by age group (same participant numbers as in H, but subset for only the genes showing higher expression in young vs old). Each point is one individual; boxes show median and IQR, red crossbar the group mean. Spearman correlation, ρ ≈ −0.14, p = 0.0105. (J) Heatmap of the same bivalent-gene signature comparing primary tumor versus adjacent normal tissue from Marteau et al^30^, showing per-gene mean standardized expression collapsed by tissue type. Color scale as in (H).

Next, we validated these findings in a recently published single cell RNA sequencing atlas of human CRC that combines 76 datasets from 48 papers^30^. This analysis showed that *TACSTD2* is repressed in epithelial cells from old versus young CRCs and adjacent normal colon (Figure 5F-G), and that a subset of the tumor bivalent gene signature is enriched in young versus old CRCs (Figure 5H-I) and in primary tumors versus adjacent normal tissue (Figure 5J). These data suggest that the epigenetically regulated developmental gene program uncovered in our mouse model is recapitulated in human disease where it is associated with the distinct biology of early-onset colorectal cancer.

Consistent with these analyses, CRISPR-engineered *Apc-*null colon organoids from young humans exhibited increased organoid size compared to CRISPR-engineered *Apc-*null colon organoids from old humans (Extended Data Figure 9A). Analysis of human adenomas by immunohistochemistry likewise revealed higher expression of TROP2 in adenomas from young patients versus old patients (Extended Data Figure 9B).

## Discussion

The incidence of CRC in younger individuals has increased significantly in this century, yet the biological factors that distinguish tumor biology in young versus aged tissue remain incompletely understood^1^. Here, we demonstrate that aging of the colon imposes an epigenetically constrained state that attenuates tumor growth following equivalent oncogenic insult. Using a focal, temporally controlled *Apc* deletion model, we show that tumors arising in aged animals are intrinsically less proliferative than those in young counterparts, revealing a growth-restrictive effect of normal aging on colorectal tumorigenesis. We propose that age-associated repression of developmental programs may represent a broader principle by which aging constrains oncogenic plasticity.

Our integrated epigenomic and transcriptomic analyses indicate that this age-associated attenuation of tumor growth is linked to progressive DNA methylation of promoters that are marked by polycomb-dependent modifications in young normal colonic/intestinal crypts and are bivalent in young tumors, bearing both H3K4me3 and H3K27me3 marks. We find that during aging, loss of histone marks signifying chromatin-dependent transcriptional states — accompanied by acquisition of DNA methylation and erosion of Polycomb-associated repression—locks these genes into a stably repressed state. Our data mirror published work implicating loss of Polycomb-associated repression accompanied by acquisition of DNA methylation^31–34^ and extend these findings by suggesting that a consequence of this aging-related biology is alteration of early tumor growth. Importantly, the epigenetic changes described here precede tumor formation and are shared between normal colonic and tumor tissue, indicating that aging remodels the epigenetic landscape upon which oncogenic mutations act, rather than being a consequence of tumor evolution.

Our results show that tumors in young animals uniquely exploit this tumor bivalent gene set that transition from Polycomb-repressed to DNA methylation-based repression with age, exhibiting increased chromatin accessibility, acquisition of H3K4me3 and transcriptional activation of developmental programs that are normally silenced with age. Many of these genes are expressed in fetal intestine, consistent with the emerging concept that tumor initiation in young tissue involves reactivation of latent developmental pathways^35^. In contrast, age-associated promoter hypermethylation restricts access to these programs, limiting tumor growth even in the presence of an oncogenic driver. These findings support a model in which aging reduces epigenetic plasticity, thereby constraining malignant potential.

Functional studies support a mechanistic role of these findings in aging-associated tumorigenesis. Pharmacologic inhibition of DNA methylation in organoids derived from aged tumors reactivated a substantial fraction of this gene set, demonstrating that DNA methylation underlies the age-associated silencing of this gene set during tumorigenesis. Although Dnmt1 inhibition may exert pleiotropic effects^36^, the preferential reactivation of genes that switch from Polycomb-dependent mechanisms to DNA methylation to maintain a repressed transcriptional state with age supports a direct contribution of DNA methylation to their repression. Conversely, genetic disruption of *Tacstd2*, a representative gene within this class, impaired organoid growth and altered expression of developmental and Wnt-associated transcriptional programs. These results suggest that sustained expression of fetal-like genes such as *Tacstd2* contributes directly to tumorigenic capacity in young tissue, whereas their epigenetic silencing with age limits tumor expansion.

Our findings have broader implications for understanding the seemingly paradoxical relationship between aging and cancer. Aging is a major risk factor for CRC, driven by the lifelong accumulation of somatic mutations and declining immune function^37^, while the incidence of early-onset CRC is rapidly rising^38^, with more aggressive and less differentiated tumors seen in younger individuals^39^. However, our data indicate that aging imposes a profound epithelial-intrinsic epigenetic barrier that actively restricts growth at early stages of tumor development by locking down critical developmental gene programs, a finding that is recapitulated in our analysis of human CRC data. This insight may help explain why early-onset colorectal cancers are diagnosed at more advanced stage with adverse histology^40^.

*TACSTD2* encodes TROP2, a developmentally and epigenetically regulated epithelial surface glycoprotein that is amplified in epithelial cancers including CRC^41^. The observation that lower *TACSTD2* expression is associated with improved survival in human colorectal cancer suggests that age-associated epigenetic states influence clinically relevant tumor phenotypes. *TACSTD2*/TROP2-targeted antibody-drug conjugates (ADCs) have been developed for treatment of several epithelial cancer types^42^. The age-dependent regulation of *TACSTD2* identified here raises the possibility that epigenetic age of the tumor may inform the utility of such targeted therapies.

Together, this work establishes that normal colonic aging is not merely a backdrop to cancer risk but an active epigenetic force that constrains oncogenic potential by stably silencing developmental gene networks. Counterintuitively, aging — long recognized as the dominant risk factor for cancer — appears protective at early stages of colorectal tumor development, providing a plausible mechanistic explanation for the distinct and aggressive biology of early-onset CRC. This mechanistic concept provides a framework for understanding how tissue aging shapes early tumor development.

## Methods

### Genetically engineered mouse model of colon tumorigenesis

The study utilized young (2-3 months) and old (18-22 months) *Apcfl/fl Villin^CreERT2^* mice^43,44^ on a pure C57BL/6 background as well as young and old C57BL/6 mice (Jackson Laboratory, # 000664). Mice were fed standard chow ad libitum. To induce tumor formation in *Apcfl/fl;Villin^CreERT^*^2^ mice, 70 µL of 100 µM 4-hydroxytamoxifen (Sigma-Aldrich,#H6278) was injected directly into the colonic submucosa using a custom 16-inch long, 45-degree bevel, 33-gauge needle (Hamilton), as previously described^45,46^. Tumor development was monitored with colonoscopy as previously described using a Karl Storz Coloview system^47–49^. Four weeks post-injection, tumors, along with normal colon crypts and small intestinal (SI) crypts, were collected for analysis. Tumors were weighed, imaged using a stereomicroscope, and processed for histological analysis. Animal experiments complied with the NIH Guide for the Care and Use of Laboratory Animals and approved by the Institutional Animal Care and Use Committee at Duke University.

### Mouse intestinal and colonic crypt isolation

Following previously published protocols, we isolated mouse intestinal and colonic crypts^50^. Briefly, the colon and SI were removed, flushed with ice-cold Ca²⁺/Mg²-free Dulbecco’s phosphate-buffered saline (DPBS) (Cytiva, #SH30378.02), opened longitudinally, and cut into 3–5 mm fragments. These fragments were rinsed three times with ice-cold DPBS and then incubated with 10 mM EDTA (EMD Millipore, #324506) in DPBS at 4°C (20 minutes for SI, 45 minutes for colon) with gentle shaking. For the SI, the tissue fragments were vigorously shaken to release crypts, then filtered through a 70-μm corning® Cell Strainer to collect the crypts. For the colon, crypts were collected directly without filtration.

### Organoid culture and growth assessment of young and old EpCAM⁺ Apc-null cells

EpCAM⁺ cells were FACS-sorted from colonic tumors of young and old Apc-null mice. Cells were embedded in 8 μL Matrigel domes at a density of 2,000 cells per dome and cultured in APC medium (Advanced DMEM/F12 (Gibco, #12634010) supplemented with 1% L-glutamine (Gibco, #35050061), 2% B27 supplement (Gibco, #17504044), 1% Antibiotic-Antimycotic solution (Gibco, #15240062), EGF (Peprotech, #31509), and 5% FBS for four days. Organoid growth was monitored via brightfield imaging using a Leica microscope at 10× magnification. The perimeter of intact, spherical organoids was manually traced and quantified using Fiji (ImageJ), and relative clonogenicity was determined by dividing the total number of organoids formed after 4 days by the initial number of cells seeded. Primary organoids were subsequently passaged, and secondary organoids were cultured under identical conditions. Organoid size was reanalyzed following secondary passage to assess self-renewal and growth potential. At least six domes per condition were analyzed in each experiment.

### EpCAM^+^ tumor epithelial cell isolation

For tumor epithelial cell isolation, tumors were chopped into small pieces and digested with 20 mg/mL of collagenase type1 (6900 U/mL) (Worthington, #LS004196) in 1X Minimal essential media for Suspension cultures (S-MEM) (Gibco, #11380-037) at 37°C for 30 - 60 mins with intermittent gentle shaking. The digestion mixture was sequentially filtered through 100-µm and 70-µm corning® Cell Strainer to obtain a single-cell suspension. For the Flow cytometer, cells were resuspended in S-MEM media and stained with fluorescent-conjugated antibodies against EpCAM (eBioscience, #12-0311-82), Ter-119-PE (BioLegend, #116208), CD45-PE (BioLegend, #103208), CD31-PE (BioLegened, #102418) for 30 min at 4°C. The Viability dye 7-AAD (7-Aminoactinomycin D) (Invitrogen, #A1310) was used to exclude dead cells. Stained cells were washed with cold PBS or FACS buffer (PBS + 1% BSA) and sorted using the following gating strategy: EpCAM^+^ (tumor epithelial cells) and CD45^-^, TER119^-^, CD31^-^ (negative selection), with 7-AAD used for viability gating. Sorted cells are further used to analyze for bulk RNA sequencing and in vitro growth study.

### Immunohistochemistry in Apc mouse tumor

Tissue samples were fixed in 10% neutral-buffered formalin (VWR # 16004-128), paraffin-embedded, and sectioned as previously described^50^. Sections were deparaffinized with xylene and graded ethanol washes, then rehydrated in water. Antigen retrieval was performed using Borg Decloaker RTU solution (Biocare Medical, #BD1000G1) in a pressure cooker (Instant Pot) for 3 minutes. Endogenous peroxidase activity was blocked using Dako Dual Endogenous Enzyme Block (Dako, #S2003), followed by incubation with 5% donkey serum (Sigma-Aldrich, #D9663) and avidin/biotin blocking (Vector Laboratories, #SP-2001) to reduce non-specific binding. Slides were incubated with a rabbit monoclonal anti-TROP2 antibody (Abcam, #ab214488), followed by detection using biotin-conjugated donkey anti-rabbit secondary antibodies (Jackson ImmunoResearch, #111-035-144). Signal amplification was achieved using the VECTASTAIN Elite ABC immunoperoxidase kit (Vector Laboratories, #PK-6100), and staining was visualized with DAB substrate (Cell Signaling Technology, #8059). Slides were counterstained with hematoxylin (Sigma-Aldrich, #GHS116), treated with bluing solution (Avantik, #RS4363-B), dehydrated, and mounted with Cytoseal (PHC Holdings Corporation, #8312-4) according to standard protocols.

### CRISPR–Cas9 knockout of *Tacstd2* in Apc-null organoids

The sorted EpCAM^+^ *Apc*-null tumor cells were cultured in 15 μL droplets of Matrigel on 6-well plates (Gen Clone, #25-105) and maintained in APC media composed of (1X Advanced DMEM F/12(Gibco, #12634-010), 2X Antibiotic-Antimycotic (Gibco, #15240062),1X GlutaMAX (Gibco, #35050-061), 50ng/mL Recombinant murine EGF (PeproTech, #315-09), 1X B27 (Thermo Fisher Scientific, #17504044), 5% Fetal Bovine Serum (Cytiva, #SH30910.03HI) and 10 μM Y-27632 dihydrochloride (APExBIO, #A3008) for 3 days. TACSTD2 sgRNA CRISPR/Cas9 plasmid (Applied Biological Materials Inc., Cat. No. 46059134) was electroporated into cells with a NEPA21 system under conditions described by Fujii et al.^51^. Total RNA was isolated according to the manufacturer’s guidelines with the following modifications: the aqueous phase containing total RNA was purified using the RNeasy Plus kit (QIAGEN, #74134). RNA was converted to complementary DNA (cDNA) with a cDNA synthesis kit (Bio-Rad, #1708891). qRT-PCR was performed with Power SYBR green master mix (Applied Biosystems, #4367659) on the Applied Biosystems Step One RT-PCR system. The following primers were used: mouse *Tacstd2* (forward: 5’-CCACCAACAAGATGACGGTCTG-3’; Reverse: 5’-AGGCACTTGGAAGTTAGCGTGG-3’), mouse HPRT (forward: 5’-AAGCTTGCTGGTGAAAAGGA-3’; Reverse: 5’-TTGCGCTCATCTTAGGCTTT-3’).

### TROP2 overexpression in aged *Apc*-mutant organoids

Lentiviral particles were produced in HEK293T cells obtained from the Duke Cell Culture Facility by co-transfecting the pLCP-Mm empty-vector control or pLCP-Trop2-Mm transfer plasmid together with the packaging plasmids psPAX2 and pMD2.G at a 5:4:2 ratio, using TransIT-293 transfection reagent (Mirus Bio) according to the manufacturer’s instructions. Viral supernatants were harvested at 48 and 72 h after transfection, pooled, cleared by centrifugation at 500 × g for 5 min, filtered through a 0.45-µm PES filter, and concentrated using Lenti-X Concentrator (Takara Bio) according to the manufacturer’s protocol. Concentrated virus was aliquoted and stored at −80 °C until use. For transduction, aged *Apc*-mutant intestinal organoids were dissociated and seeded onto plates pre-coated with 1% Matrigel in organoid growth medium supplemented with concentrated lentivirus and 8 µg ml⁻¹ polybrene. After 48 h, transduced organoids were re-embedded in Matrigel domes, and selection was performed with 1 µg ml⁻¹ puromycin (Gibco) for 5 days, with medium refreshed every 48 h. Trop2 overexpression was confirmed by RT–qPCR and immunoblotting.

### Immunoblotting

Organoids were harvested with Matrigel domes and incubated in Cell Recovery Solution to dissolve Matrigel, followed by two washes with culture medium. Organoid lysates were prepared in RIPA buffer (Sigma, #SLCD5849) supplemented with Halt™ Protease and Phosphatase Single-Use Inhibitor Cocktail (Thermo Scientific, #ZH397539). Protein concentrations were determined using a Pierce BCA Protein Assay Kit (# VA294769A). Equal amounts of protein lysate, typically 20–25 μg per sample, were resolved by SDS-PAGE and transferred onto nitrocellulose membranes (Pall Corporation, USA). Membranes were incubated with antibodies against TROP2/TACSTD2 (Abcam, #ab214488) and β-actin (Cell Signaling Technology, #4970S), followed by HRP-conjugated secondary antibody (Cell Signaling Technology, #7074S). Bands were detected using SuperSignal™ chemiluminescent substrate (Thermo Scientific, #186394) and imaged with a Bio-Rad ChemiDoc Imaging System.

### Culture of AKPST cancer organoids

AKPST organoids were dissociated into single cells and combined with Matrigel (Corning, #356231) at a 1:3 ratio. Cell-Matrigel suspensions were plated as 20 µL domes (7–8 domes per well) in a 6-well plate, followed by polymerization at 37°C for 10–15 minutes. Organoids were cultured in Advanced DMEM/F-12 medium (Gibco, #12634010) supplemented with 5% fetal bovine serum (Cytiva, #SH3091003), 2% B27 supplement (Gibco, #17504044), 1% L-glutamine (Gibco, #35050061), and 1% Antibiotic-Antimycotic solution (Gibco, #15240062) at 37°C in a 5% CO₂ incubator. The media was refreshed every 2–3 days. For passaging, Matrigel domes were dissolved with cell recovery solution (Corning, #354270), and organoids were enzymatically dissociated with TrypLE Express (Gibco, #12605010) to generate a single-cell suspension before re-plating.

### Human organoid culture and APC editing

Human colon tissue samples from young (29–48 years) and aged (72–77 years) individuals were obtained from endoscopic biopsies collected at Duke University Hospital under an approved Duke Institutional Review Board protocol (IRB #00104775). Colonic crypts were isolated using 10 mM EDTA chelation and subsequently embedded in growth factor–reduced Matrigel (Corning # 356231) for three-dimensional culture. Organoids were maintained in complete medium consisting of 50% L-WRN conditioned medium supplemented with 1× B27 (Gibco) and 1× N2 (Gibco), along with the following growth factors and small molecules: 50 ng/mL epidermal growth factor (EGF; R&D Systems), 10 μM SB202190 (Sigma), 500 nM A83-01 (Tocris Bioscience), 10 nM gastrin I (Sigma), 10 mM nicotinamide (Sigma), 1 mM N-acetylcysteine (Sigma), and 100 μg/mL Primocin (InvivoGen). To enhance cell survival following crypt isolation and passage, 10 μM Y-27632 (a ROCK inhibitor) was included in the culture medium for the first 48 hours to prevent anoikis, as previously described^52^.

Targeted disruption of the APC gene in human colon organoids was performed using CRISPR/Cas9 genome editing, as described^53^. Briefly, organoids were enzymatically dissociated into small multicellular fragments (approximately 5–10 cells per fragment) and electroporated with ribonucleoprotein (RNP) complexes consisting of recombinant Cas9 protein and synthetic single-guide RNAs (sgRNAs) targeting APC. Electroporation was carried out using the NEPA21 electroporator (Nepagene) according to the manufacturer’s recommendations. Following electroporation, cells were embedded in Matrigel and cultured in complete medium to allow recovery and organoid formation over 7 days. To enrich for APC-mutant clones, organoids were subsequently cultured in medium lacking Wnt3a and R-spondin. Under these selective conditions, control organoids treated with non-targeting sgRNAs ceased proliferation within 3–5 days, whereas APC-targeted organoids exhibited sustained growth and expansion. Genome editing efficiency was initially assessed using the SURVEYOR nuclease assay (Transgenomic), and candidate clones were further validated by Sanger sequencing to confirm indel formation at the APC locus.

### *Apc*-null organoid treatment with Dnmt1 inhibitor

The isolated EpCAM^+^ *Apc*-null tumor cells from young and old mice were collected in APC media. Cells were then plated on growth factor-reduced Matrigel domes (Corning, #356231) and cultured in APC media. To inhibit anoikis, the media were supplemented with 10 µM Y-27632 dihydrochloride (APExBIO, #A3008). For Dnmt1 inhibition, cells were treated with 2 µM of GSK-3484862 (Medchemexpress, #HY-135146) or DMSO for 4 days. Following treatment, organoids were washed with 1× PBS, collected in TRIzol reagent (Thermo Scientific, #15596026), and stored at –80 °C for downstream RNA analysis.

### Human colonic polyp samples and immunohistochemistry

Human colonic polyp specimens were obtained from the Nashville Veterans Affairs Tissue Biorepository under IRB protocol #1705818 and used in accordance with all relevant Department of Veterans Affairs guidelines and procedures. Early-onset polyps were defined as specimens from individuals younger than 45 years at the time of colonoscopy, whereas typical age-onset polyps were obtained from individuals aged 60–65 years at colonoscopy. All tissues were collected as part of routine clinical care between 2006 and 2015. Formalin-fixed, paraffin-embedded tissues were sectioned at 5 μm thickness using a Jung Histocut microtome.

Immunohistochemical staining was performed as previously described, with antigen retrieval in pH 6.0 citrate buffer^54^. Sections were incubated with anti-human TROP-2 antibody (R&D Systems, #AF650), followed by ImmPRESS-HRP Anti-Goat IgG detection reagent (Vector Laboratories) according to the manufacturer’s instructions.

### DNA methylation array

DNA was extracted from normal colon crypts and *Apc*-null tumors with the AllPrep DNA/RNA kit (Qiagen, #80204). Using 1 µg of starting DNA, bisulfite conversion was performed using the EZ DNA Methylation-Lightning Automation kit (Zymo Research, #D5049). DNA methylation was detected with ∼200 ng of bisulfite converted DNA using Infinium MouseMethylation-12 v1.0 BeadChip arrays (Illumina, # 20041558) following the Illumina Infinium HD methylation protocol.

DNA methylation array data was analyzed with R package SeSAMe (version 1.21.7)^55^. The differential methylated positions (DMPs) were identified with p value < 0.0001 and absolute estimated methylation level > 0.2. Adjusted p value threshold 0.001 has been applied to identify the differential methylated regions (DMRs). We performed the GO terms enrichment based on default settings for gene sets associated with DMRs by GREAT analysis software tool^56^.

### Cleavage under targets and tagmentation (CUT&Tag)

CUT&Tag was performed as reported with minor modifications^18^. Briefly, nuclei were extracted from snap-frozen tissue and cells with single nucleus isolation kit (Invent Biotechnology, #SN-047) following the manufacturer’s protocol. After lightly fixing with 0.1% formaldehyde for 2 min at room temperature, 20,000 - 50,000 nuclei were used. Antibodies against H3K4me3 (Active Motif, #39159), and H3K27me3 (Active Motif, #39055) were used as primary antibodies at 1:100 dilution. Tagmented DNA was purified with Qiagen MinElute PCR purification kit (QIAGEN, #28004) and amplified with High-fidelity PCR mix (NEB, #M0541S). Libraries were purified with AMPure XP beads (Beckman, #A63881) and sequenced on Illumina NextSeq 500 or NovaSeq 6000.

### ATAC-seq

Nuclei were isolated from snap-frozen tissues and cells and slightly fixed as described for CUT&Tag. 50,000-100,000 nuclei were incubated with 30 µl of tagmentation buffer (Illumina, # 20034211) containing 15 µl of 2X Tagment DNA buffer, 3 µl of Tn5, and 12 µl of H2O, at 37°C for 30 min. Tagmented DNA was purified, amplified, and purified as described for CUT&Tag. Libraries were sequenced on Illumina NextSeq 500 or NovaSeq 6000 instruments.

### Data analysis for CUT&Tag and ATAC-seq

Raw read pairs were mapped to the mm10 reference genome by Bowtie v2.1^57^. Alignments were filtered to retain only proper pairs with MAPQ at least 5 via samtools (-q 5-f 2). Duplicate fragments were removed with MarkDuplicates.jar from the Picard tool suite v1.1. Peak calls for histone modifications and ATAC-seq were made by HOMER v4.10 and MACS2, respectively, followed by filtering out calls with overlap to mm10 blacklist regions^58,59^. For each histone mark and ATAC-seq, a set of unified peaks was defined as regions called as peaks in 2 out of 5 samples; unified peaks are limited to canonical chromosomes, neighboring regions within 50bp are merged, and a minimum peak size of 50 bp is applied. Coverage tracks were generated by converting paired-end data to a single fragment (bedtools bamtobed with “-bedpe” option then extracting columns for chromosome and first/last coordinate), converting to bedGraph (bedtools genomecov with “-bg” option), scaled to 3 million fragments per sample, then converted to bigWig format via UCSC utility bedGraphToBigWig. Differential peaks were identified with EdgeR v3.34.1 at FDR 0.005 for H3K27me3 and ATAC-seq^60^. Peaks annotation was conducted by R package ChIPseeker^61^. Motif analysis was carried out by using the HOMER v4.10 with unchanged peaks as background^59^. Correlation of alterations between H3K27me3 enrichment and DNA methylation of DMRs during aging was calculated with normalized read counts mapped to DMRs, excluding the DMRs with low number (<50) of total mapped fragment count, and estimated methylation level difference of DMRs. Gene ontology analysis was conducted using DAVID web service^62^. To assess the statistical significance of overlap between two sets of intervals, we conducted Fisher’s exact test using the BEDTools fisher function (‘bedtools fisher-a a.bed-b b.bed-g mm10.chromSizè)^63^. For calculating the extremely small P-values, we used the hypergeometric distribution function phyper in R with ‘log.p =TRUE’ option.

### RNA-seq

Total RNA was extracted from fresh/snap-frozen tissue with TRIzol Reagent (Thermo Scientific, #15596026). After rRNA depletion with Ribo-Zero H/M/R Gold rRNA Removal Kit (Illumina, #MRZG126), purified RNA was used for RNA-seq library preparation with TruSeq Stranded Total RNA Library Prep Kit. RNA-seq libraries were sequenced on Illumina NextSeq 500.

Sequencing reads were mapped to mm10 reference genome with STAR v2.7.9a at default parameters^64^, and then differentially expressed genes (DEGs) were identified with FDR < 0.05 cutoff by R package edgeR (version 3.34.1)^60^.

### DNA methylation in colon adenocarcinoma (COAD) patients

DNA methylation of gene promoters in COAD patients was analyzed with UALCAN platform^65^. COAD patients in TCGA dataset were grouped by age and significance of difference between different groups was estimated by student’s t-test.

### Kaplan-Meier survival curve

Kaplan–Meier RNA-seq survival analysis of the overall survival probability for colon cancer patients with low and high *TACSTD2* expression was conducted by the KM plotter^66^.

### Analysis of Human CRC bulk RNAseq data

We queried TCGA-COAD^27^ data using “*TCGAbiolink*” package in R (v.4.5.2). Bulk RNAseq data of tumors comprised a total of 477 samples. We used raw gene read counts and DESeq2 with default parameters to compute fold changes and differences between EOCRC and LOCRC tumors normalized to control samples. We generated mean *z*-score per sample (fetal-like and bivalent signature scores) from the log2 fold changes of the gene expression. The fetal-like gene signature has been previously described by Mustata et al^28^. We queried the human orthologs of these genes using the *gprofiler* package in R (v.4.5.2).

Heat maps of gene expression changes were generated using *pheatmap* package in R. The significance of differences between EOCRC and LOCRC groups was determined by unpaired nonparametric Wilcoxon test.

### Analysis of Human CRC single-cell RNAseq data

#### Data Analysis of Early Onset CRC/Late Onset CRC human data

Data from a previously published atlas^30^ of human colorectal cancer single-cell RNA sequencing studies were reanalyzed in RStudio using Seurat package v5^67^.

#### Cell type annotation and filtering

Single-cell transcriptomes were annotated using a previously established atlas cell type reference^30^. Data were subset to only include participants with age data. Cancer cells and epithelial cells were identified and isolated based on “atlas_cell_type_coarse” annotation, and downstream analyses were restricted to these populations (634,535 cells, 359 individuals).

#### Age grouping and aggregation

Participants were stratified into age groups using custom split binning scheme with divisions at 45 and 70 years old. For each individual and age group combination, gene expression was aggregated by calculating the mean expression per person. To avoid pseudoreplication arising from the nested cells-within-participants structure of the data, per-cell values were aggregated to the individual level: for each person within each tissue compartment, the mean value across that participant’s epithelial cells was computed, and individuals contributing fewer than 20 cells in each compartment were excluded. Each data point therefore represents a single individual.

#### Module scoring

Gene set module scores were computed using the Seurat *AddModuleScore* function^67^ for a curated set of human bivalent genes (n=45 genes). Module scores represent the average normalized expression of genes in each set, adjusted for background expression of an expression-bin-matched, randomly selected control gene set (default parameters). Scores were computed for curated human bivalent (n = 45 genes) gene sets, as well as for three subsets of the bivalent genes: a set of genes that are were upregulated in young-vs-old tumors (n = 23), a set that were upregulated in primary tumor-vs-adjacent normal (n = 34), and the overlap of these two (n = 20).

#### Statistical analysis

Associations between age groups and module scores were assessed separately for each tissue type (primary tumor, adjacent normal). Monotonic age-related trend assessed by Spearman rank correlation, treating age group as an ordinal variable. Statistical tests were conducted using base R functions, with p-values calculated from per-participant aggregated data (minimum 20 cells per participant).

#### Visualization

Per-participant values were displayed as box-and-whisker plots (box = interquartile range, whiskers = 1.5×IQR, black center line = median, red line = group mean) with individuals overlaid as jittered points, generated using ggplot2.

#### TACSTD2 per-participant expression by age

Participant-matched cancer epithelial and adjacent normal cells were isolated from integrated atlas based on coarse cell-type annotation. Log-normalized TACSTD2 expression was extracted from the RNA data layer, and for each participant the mean log-normalized expression was computed across all that participant’s cancer cells, giving one value per individual. Participants were stratified into three age groups — young (<45), middle (45–70), and old (>70) — and primary tumor and adjacent normal tissue were analyzed separately. Per-person values were shown as box-and-whisker plots (box = interquartile range, whiskers = 1.5×IQR, black center line = median, red line = mean) with individuals overlaid as jittered points. Monotonic trend with age was assessed by Spearman rank correlation (age group treated as ordinal). Analyses were performed in RStudio using Seurat^67^, dplyr^68^, ggplot2^69^, and ggpubr^70^.

## Acknowledgements

We gratefully acknowledge the Genomics Core Facility at NIEHS for assistance with NGS library sequencing. We thank Laura Wharey for her help with the DNA methylation microarray experiment. We thank Omer H. Yilmaz for sharing AKPST organoids. We are grateful to Zachary C. Hartman for generously sharing the pLCP-Mm and pLCP-Trop2-Mm plasmids used in this study. This manuscript was improved by critical reading from Dr. Raja Jothi, Dr. Xiaoling Li, Dr. Jackson Hoffman, Dr. Joseph Rodriguez, Dr. Harriet Kinyamu and members of the Wade and Roper laboratories.

This research was supported in part by the Intramural Research Program of the NIH, National Institute of Environmental Health Sciences. The contributions of the NIH authors are considered Works of the United States Government. The findings and conclusions presented in this paper are those of the authors and do not necessarily reflect the views of the NIH or the U.S. Department of Health and Human Services.

## Data availability

All DNA methylation data in this study has been uploaded to Gene Expression Omnibus under the accession GSE271396. The RNA-seq, ATAC-seq. and CUT&Tag datasets have been deposited in the Gene Expression Omnibus repository, under accession numbers: GSE272054, GSE332724 and GSE284524.

