## Extended Data for "Aging restricts colorectal tumor growth by epigenetically silencing developmental gene programs"

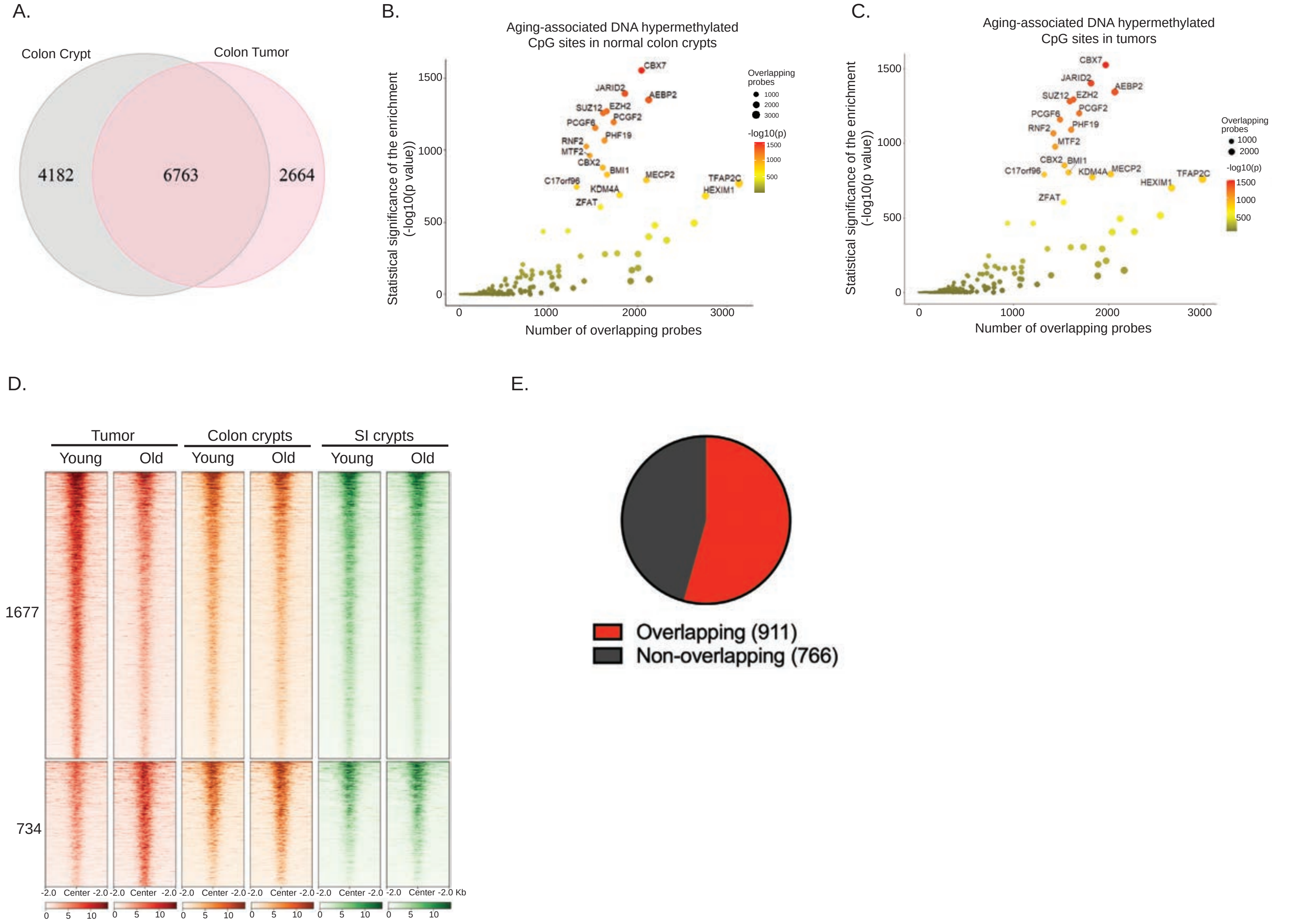


**Extended Data Figure 1. DNA methylation and chromatin accessibility changes during aging**

(A) Overlapped DMPs between tumors and matched normal crypts during aging.
(B-C) Association of aging-related hypermethylated CpG sites in normal colonic crypts (B) and tumors (C) with the occupancy of transcriptional regulators. x-axis denotes the number of aging-associated hypermethylated probes overlapping binding sites, and y-axis represents statistical significance of the enrichment (Fisher's exact test).
(D) Heat map showing differential ATAC peaks defined in tumors during aging.
(E) Pie chart showing reduced ATAC peaks overlapped with aging-related hypermethylated DMRs.

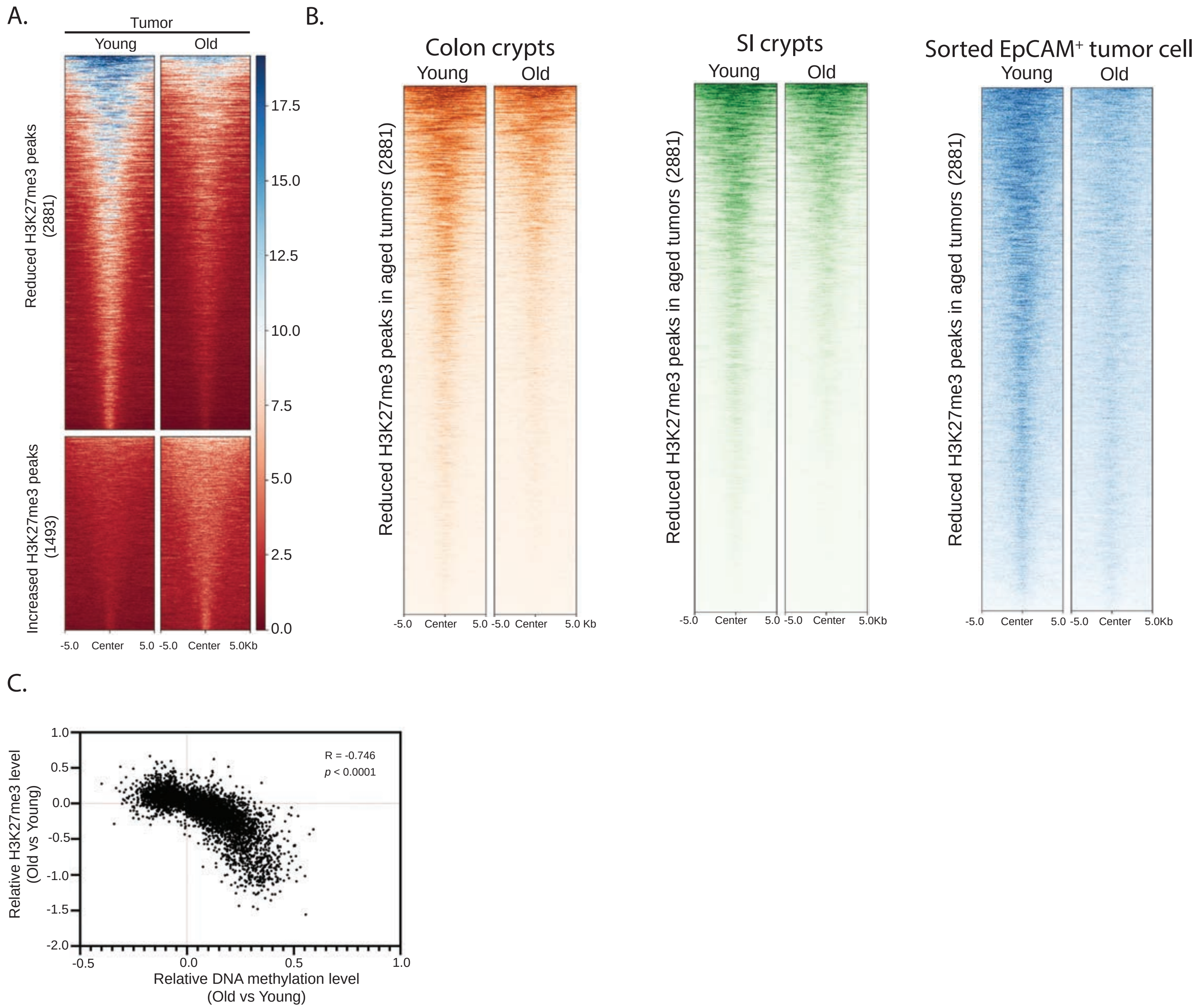


**Extended Data Figure 2. Alterations of H3K27me3 histone modification during aging**

(A) Heat map displaying the differentially enriched H3K27me3 peaks between young and old colorectal tumors.
(B) Heatmap and metaplot showing the H3K27me3 levels for 2881 peaks, which exhibiting lower occupancy of H3K27me3 in old tumors, in young and old normal colonic crypts, SI crypts, and sorted EpCAM+ tumor cells.
(C) Scatter plot showing the correlation between H3K27me3 enrichment and DNA methylation levels for DMRs in tumors during aging.

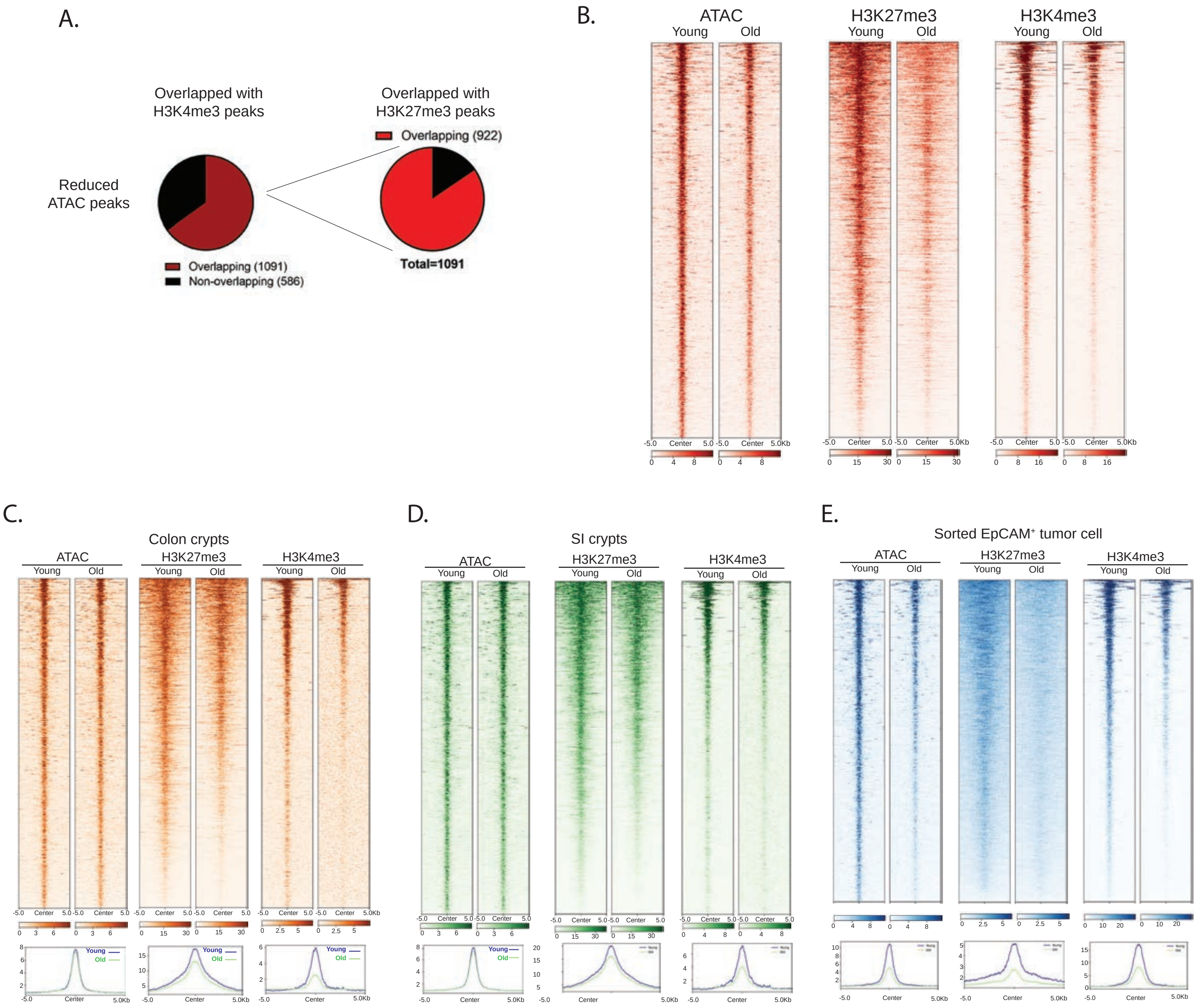


**Extended Data Figure 3. Aging-associated epigenetic alterations at tumor bivalent chromatin**

(A) Reduced ATAC peaks as aging in tumors overlapped with H3K4me3 and H3K27me3 marks.
(B) Heat map and metaplot showing the chromatin accessibility and bivalent marks (H3K4me3 and H3K27me3) for 922 bivalent domains with reduced accessibility during aging.
(C-E) Heatmap and metaplot showing epigenetic changes of 922 tumor bivalent domains in normal colonic crypts (C), SI crypts (D), and sorted EpCAM+ tumor cells (E) from young and old animals.

Yang Liu et al Extended Data Figure 4

A.

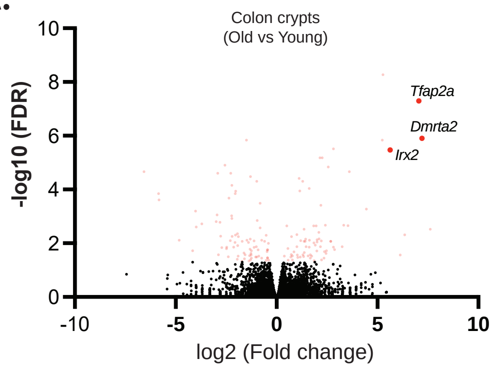


**Extended Data Figure 4. Aging-related gene expression changes in normal colon crypts**

(A) Volcano plot showing transcriptomic changes in normal colonic crypts during aging.

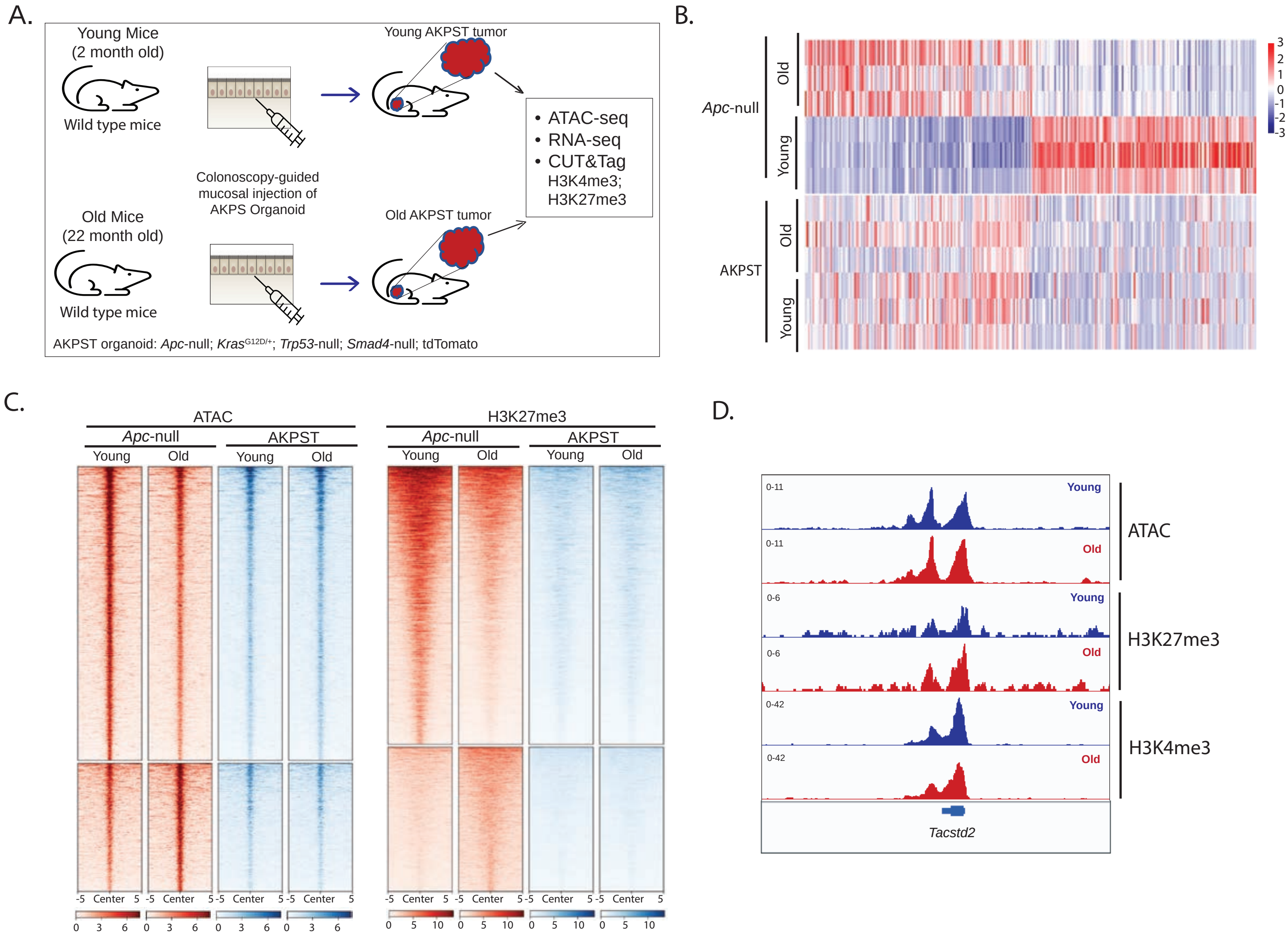


**Extended Data Figure 5. AKPST organoid-implanted CRC tumors from young and old mice.**

(A) APKST organoids were implanted into young and old wild type C57BL6/J mice via colonoscopy-guided mucosal injection for inducing CRC.
(B) Heat map showing the gene expression levels of aging-related DEGs identified in Apc-null tumors.
(C) Heat maps showing the enrichment of epigenetic marks for aging-related differential peaks detected in Apc-null CRC.
(D) IGV tracks displaying the histone modifications enrichment and chromatin accessibility at Tacstd2 gene in AKPST tumors from young and old mice.

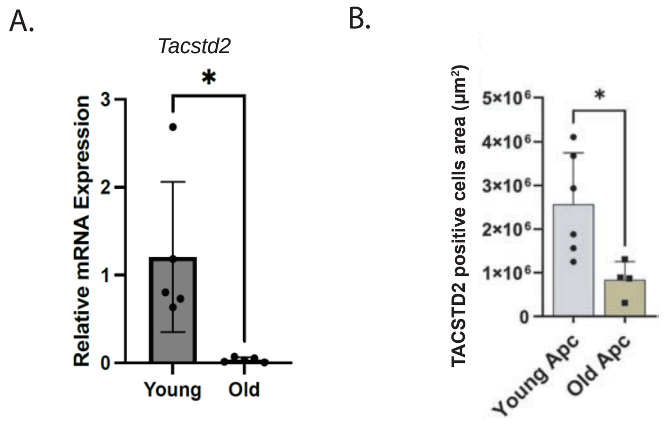


**Extended Data Figure 6. Repression of Tacstd2 expression in old Apc-null tumor**

(A) Relative Tacstd2 mRNA expression in young and old Apc-null organoids. Data are mean ± s.d. from independent biological replicates. Statistical significance was assessed by an unpaired two-tailed Student's t-test. $P < 0.05$.
(B) Quantification of TACSTD2-positive staining area in young and old Apc-null tumors by immunohistochemistry. Data are mean ± s.d. from independent biological replicates. Statistical significance was assessed by an unpaired two-tailed Student's t-test. $P < 0.05$.

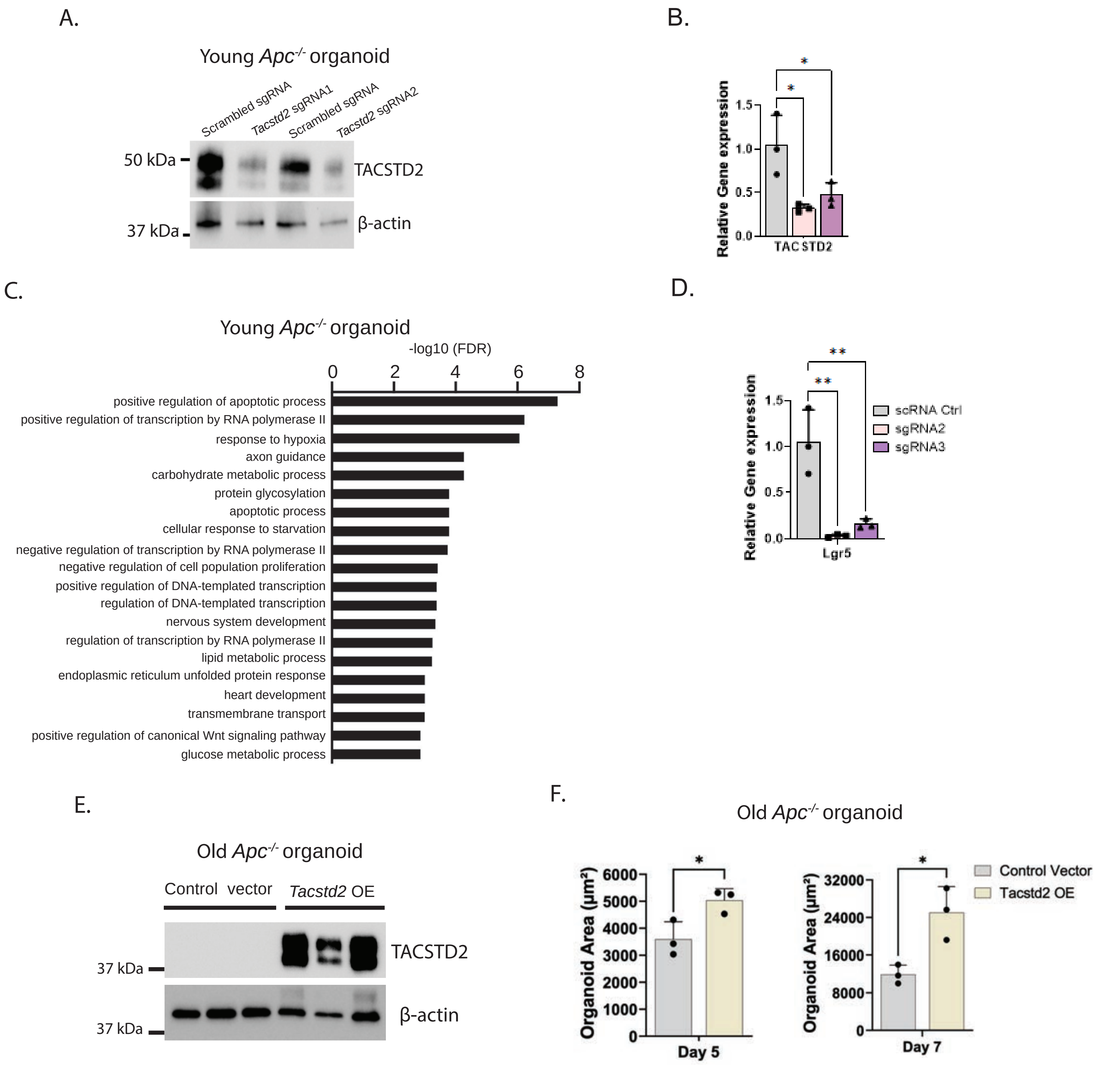


**Extended Data Figure 8. Knockout and overexpression of Tacstd2 in tumor organoid**

(A) Immunoblot showing the knockout efficiency of Tacstd2 following Cas9/sgRNA expression.
(B) RT-PCR analysis of Tacstd2 expression in organoids expressing scramble sgRNA (grey), Tacstd2 sgRNA 1 (pink) or Tacstd2 sgRNA 3 (purple).
(C) Pathways associated with down-regulated genes following Tacstd2 knockout.
(D) RT-PCR analysis of Lgr5 expression in organoids expressing scramble sgRNA (grey), Tacstd2 sgRNA 1 (pink) or Tacstd2 sgRNA 3 (purple column).
(E) Representative immunoblot analysis of Tacstd2 expression in old Apc-null organoids transduced with control vector or Tacstd2 overexpression construct. β-actin, loading control. Each lane represents an independent organoid line (n=3).
(F) Quantification of organoid area in old Apc-null organoids transduced with control vector or Tacstd2 overexpression construct at day 5 and day 7. Data are mean ± s.d. from independent biological replicates (n=3). Statistical significance was assessed by an unpaired two-tailed Student's t-test. $P < 0.05$.

A.

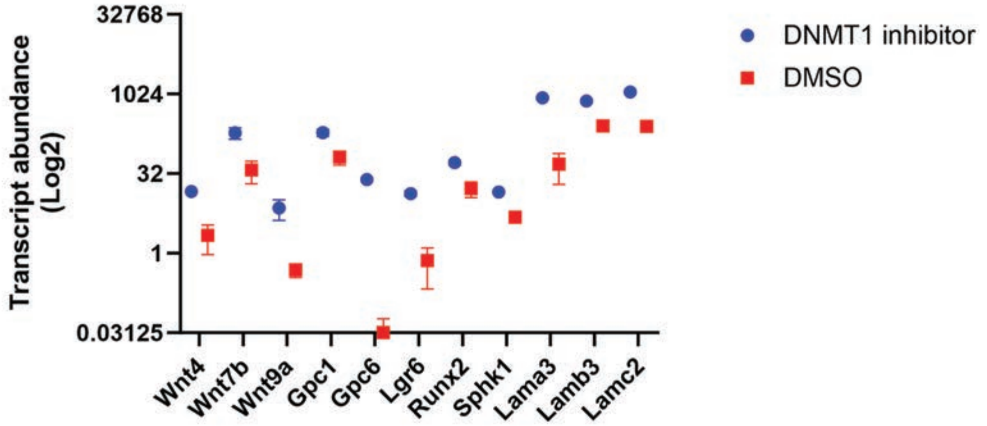


**Extended Data Figure 7. Derepression of Wnt signaling-related genes by DNMT1 inhibitor treatment in old tumor organoids**

(A) Relative gene expression of Wnt signaling-related genes in old tumor organoids treated with DMSO or DNMT1 inhibitor, GSK3484862.

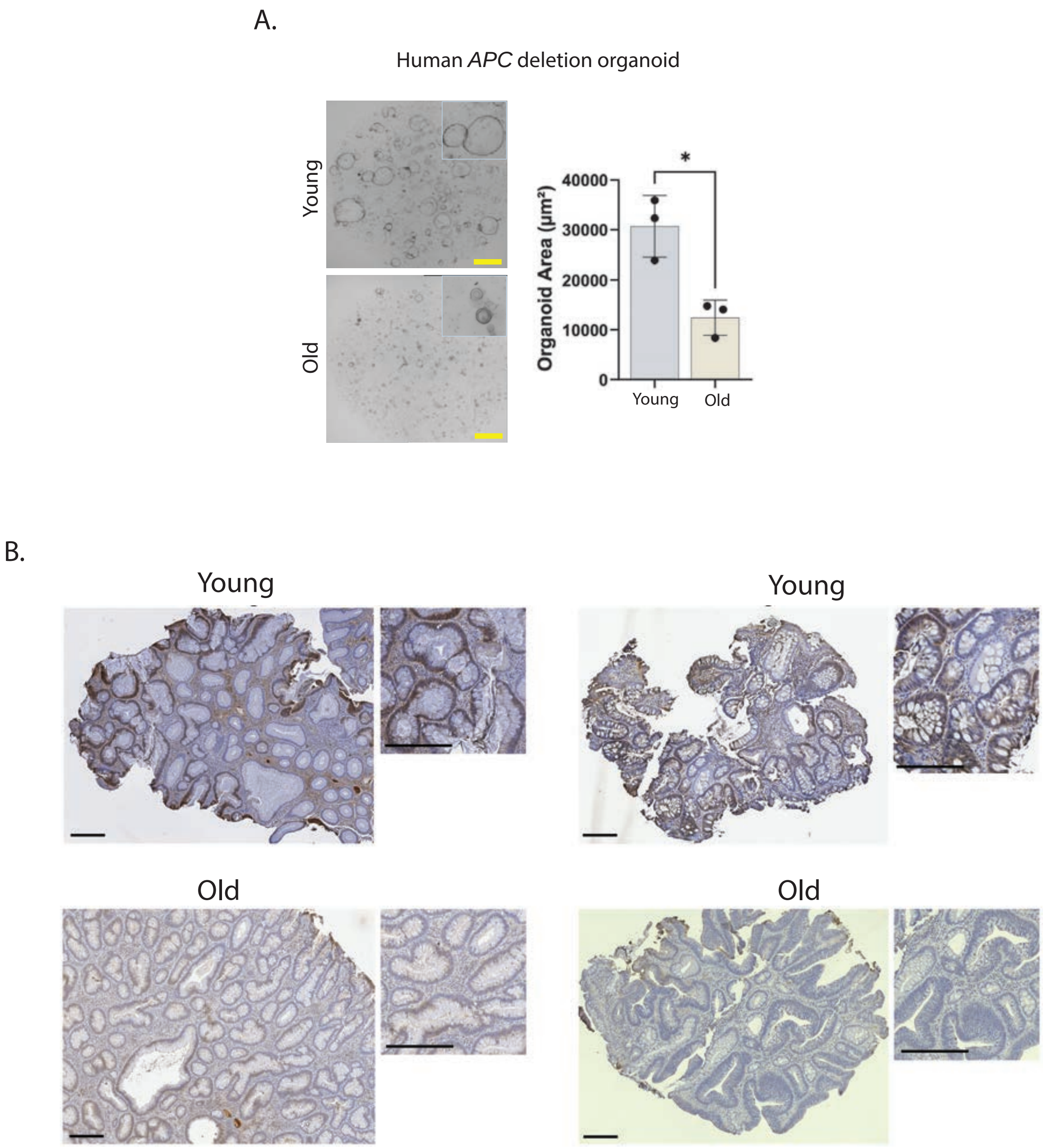


**Extended Data Figure 9. Analysis of APC deletion in human organoids and expression of TACSTD2 in human adenomas**

(A) Representative bright-field images (left) and quantification of organoid area (right) of human APC-null tumor organoids derived from young adult (29–48 years) and aged (72–77 years) donors. Insets, higher magnification. Scale bars, 100 µm. Data are mean ± s.e.m.; n = 3 independent experiments. *P < 0.05, two-tailed Student's t-test (or Mann–Whitney U-test, as appropriate).
(B) Representative microscopy images reveal immunohistochemistry staining of TACSTD2 in tubular adenomas. "Young" refers to polyps from individuals aged 42 and 44 years and "Old" refers to polyps from individuals aged 62 years. Scale bars = 100 uM.

Extended Data Table 1. The majority of 46 genes are targets of Wnt signaling.

© 2000-2026 QIAGEN. All rights reserved.

| ID | Genes in dataset | Prediction (based on measurement direction) |
|---|---|---|
| Adgrg6 | ADGRG6 | Affected |
| Cadps | CADPS | Affected |
| Chd3 | CHD3 | Affected |
| Col4a1 | COL4A1 | Affected |
| Dnajc6 | DNAJC6 | Affected |
| Eid2 | EID2 | Affected |
| Erbb4 | ERBB4 | Affected |
| F5 | F5 | Affected |
| Fras1 | FRAS1 | Affected |
| Gata5 | GATA5 | Affected |
| Ltbp2 | LTBP2 | Affected |
| Macroh2a2 | MACROH2A2 | Affected |
| Mn1 | MN1 | Affected |
| Nebl | NEBL | Affected |
| Onecut2 | ONECUT2 | Affected |
| Peg3 | PEG3 | Affected |
| Plxdc2 | PLXDC2 | Affected |
| Ppic | PPIC | Affected |
| Rarb | RARB | Affected |
| Rbp1 | RBP1 | Affected |
| Runx1t1 | RUNX1T1 | Affected |
| Sall2 | SALL2 | Affected |
| St8sia6 | ST8SIA6 | Affected |
| Tacstd2 | TACSTD2 | Affected |
| Tiam1 | TIAM1 | Affected |

The table depicts output from Ingenuity Pathway Analysis of the 46 gene set described in the text. Genes identified as Wnt signaling targets are listed (25 of 46).

Extended Data Table 2

Top 25 GO Biological Processes

| Index | Name | P-value | Adjusted p-value | Odds Ratio | Combined score |
|---|---|---|---|---|---|
| 1 | Heart Induction (GO:0003129) | 0.0001436 | 0.03209 | 151.12 | 1337.16 |
| 2 | Regulation of Heart Morphogenesis (GO:2000826) | 0.0001844 | 0.03209 | 129.53 | 1113.72 |
| 3 | Negative Regulation of Transcription by RNA Polymerase II (GO:0000122) | 0.002221 | 0.1094 | 4.33 | 26.43 |
| 4 | Positive Regulation of Cell Projection Organization (GO:0031346) | 0.002348 | 0.1094 | 12.36 | 74.83 |
| 5 | Negative Regulation of DNA-templated Transcription (GO:0045892) | 0.002844 | 0.1094 | 3.73 | 21.87 |
| 6 | Positive Regulation of Notch Signaling Pathway (GO:0045747) | 0.003102 | 0.1094 | 26.63 | 153.81 |
| 7 | Regulation of Cell Motility (GO:2000145) | 0.003110 | 0.1094 | 11.16 | 64.41 |
| 8 | Positive Regulation of Axonogenesis (GO:0050772) | 0.003633 | 0.1094 | 24.47 | 137.46 |
| 9 | Aortic Valve Morphogenesis (GO:0003180) | 0.003633 | 0.1094 | 24.47 | 137.46 |
| 10 | Regulation of Axonogenesis (GO:0050770) | 0.003633 | 0.1094 | 24.47 | 137.46 |
| 11 | Aortic Valve Development (GO:0003176) | 0.004202 | 0.1094 | 22.63 | 123.83 |
| 12 | Visual System Development (GO:0150063) | 0.004202 | 0.1094 | 22.63 | 123.83 |
| 13 | Negative Regulation of Cell Motility (GO:2000146) | 0.004255 | 0.1094 | 9.95 | 54.30 |
| 14 | Digestive Tract Development (GO:0048565) | 0.004401 | 0.1094 | 22.08 | 119.79 |
| 15 | Regulation of Cell Migration (GO:0030334) | 0.004924 | 0.1142 | 4.96 | 26.35 |
| 16 | Regulation of Stem Cell Differentiation (GO:2000736) | 0.005910 | 0.1285 | 18.85 | 96.72 |
| 17 | Negative Regulation of G Protein-Coupled Receptor Signaling Pathway (GO:0045744) | 0.007112 | 0.1294 | 17.07 | 84.42 |
| 18 | Positive Regulation of DNA-templated Transcription (GO:0045893) | 0.009070 | 0.1294 | 3.03 | 14.25 |
| 19 | Negative Regulation of Cell Migration (GO:0030336) | 0.009091 | 0.1294 | 7.50 | 35.24 |
| 20 | Positive Regulation of Neurogenesis (GO:0050769) | 0.009818 | 0.1294 | 14.35 | 66.35 |
| 21 | Eye Development (GO:0001654) | 0.01101 | 0.1294 | 13.49 | 60.84 |
| 22 | Establishment of Blood-Brain Barrier (GO:0060856) | 0.01145 | 0.1294 | 110.83 | 495.42 |

| Index | Name | P-value | Adjusted p-value | Odds Ratio | Combined score |
|---|---|---|---|---|---|
| 23 | Renal Tubule Development (GO:0061326) | 0.01145 | 0.1294 | 110.83 | 495.42 |
| 24 | Axon Choice Point Recognition (GO:0016198) | 0.01145 | 0.1294 | 110.83 | 495.42 |
| 25 | Positive Regulation of Vitamin D Receptor Signaling Pathway (GO:0070564) | 0.01145 | 0.1294 | 110.83 | 495.42 |

Top 25 MGI Mammalian Phenotypes

| Index | Name | P-value | Adjusted p-value | Odds Ratio | Combined score |
|---|---|---|---|---|---|
| 1 | Abnormal Glomerular Capsule Parietal Layer Morphology MP:0011498 | 0.0001436 | 0.03804 | 151.12 | 1337.16 |
| 2 | Postnatal Growth Retardation MP:0001732 | 0.0001575 | 0.03804 | 5.95 | 52.09 |
| 3 | Myocardial Trabeculae Hypoplasia MP:0010503 | 0.0003972 | 0.05877 | 82.41 | 645.35 |
| 4 | Abnormal Prepulse Inhibition MP:0003088 | 0.0005331 | 0.05877 | 69.72 | 525.49 |
| 5 | Decreased Circulating Interferon-Beta Level MP:0008576 | 0.0006084 | 0.05877 | 64.74 | 479.38 |
| 6 | Delayed Kidney Development MP:0000528 | 0.0009577 | 0.06608 | 50.34 | 349.94 |
| 7 | Abnormal Glomerular Mesangium Morphology MP:0011339 | 0.0009577 | 0.06608 | 50.34 | 349.94 |
| 8 | Abnormal Basement Membrane Morphology MP:0004272 | 0.001383 | 0.07745 | 41.18 | 271.12 |
| 9 | Abnormal Lung Development MP:0001176 | 0.001443 | 0.07745 | 14.74 | 96.41 |
| 10 | Abnormal Renal Glomerulus Morphology MP:0005325 | 0.002650 | 0.1158 | 11.83 | 70.19 |
| 11 | Neonatal Lethality, Incomplete Penetrance MP:0011088 | 0.004117 | 0.1158 | 6.67 | 36.62 |
| 12 | Abnormal Bone Structure MP:0003795 | 0.004168 | 0.1158 | 6.64 | 36.41 |
| 13 | Abnormal Blood Circulation MP:0002128 | 0.004202 | 0.1158 | 22.63 | 123.83 |
| 14 | Narrow Eye Opening MP:0005287 | 0.005022 | 0.1158 | 20.57 | 108.89 |
| 15 | Increased Cornea Thickness MP:0011962 | 0.005022 | 0.1158 | 20.57 | 108.89 |
| 16 | Increased Myocardial Fiber Size MP:0004564 | 0.005682 | 0.1158 | 19.25 | 99.54 |
| 17 | Decreased Trabecular Bone Volume MP:0010879 | 0.005910 | 0.1158 | 18.85 | 96.72 |
| 18 | Microphthalmia MP:0001297 | 0.007255 | 0.1158 | 5.65 | 27.81 |
| 19 | Abnormal Iris Morphology MP:0001322 | 0.007621 | 0.1158 | 16.45 | 80.20 |
| 20 | Kidney Inflammation MP:0001859 | 0.008147 | 0.1158 | 15.87 | 76.32 |

| Index | Name | P-value | Adjusted p-value | Odds Ratio | Combined score |
|---|---|---|---|---|---|
| 21 | Abnormal Kidney Morphology MP:0002135 | 0.008238 | 0.1158 | 4.36 | 20.92 |
| 22 | Glomerulosclerosis MP:0005264 | 0.008688 | 0.1158 | 15.33 | 72.74 |
| 23 | Micrognathia MP:0002639 | 0.009530 | 0.1158 | 14.58 | 67.86 |
| 24 | Abnormal Hindlimb Morphology MP:0000556 | 0.01041 | 0.1158 | 13.91 | 63.50 |
| 25 | Cataract MP:0001304 | 0.01157 | 0.1158 | 4.91 | 21.88 |

Top 25 KEGG Pathways

| Index | Name | P-value | Adjusted p-value | Odds Ratio | Combined score |
|---|---|---|---|---|---|
| 1 | ECM-RECEPTOR INTERACTION | 0.01748 | 0.3888 | 10.50 | 42.50 |
| 2 | SMALL CELL LUNG CANCER | 0.01900 | 0.3888 | 10.03 | 39.76 |
| 3 | ATP-DEPENDENT CHROMATIN REMODELING | 0.02651 | 0.3888 | 8.35 | 30.32 |
| 4 | INTEGRIN SIGNALING | 0.04833 | 0.4269 | 5.96 | 18.06 |
| 5 | CHEMOKINE SIGNALING PATHWAY | 0.07144 | 0.4269 | 4.75 | 12.54 |
| 6 | PROTEOGLYCANS IN CANCER | 0.07536 | 0.4269 | 4.61 | 11.91 |
| 7 | CYTOSKELETON IN MUSCLE CELLS | 0.09809 | 0.4269 | 3.93 | 9.13 |
| 8 | ENDOCYTOSIS | 0.1112 | 0.4269 | 3.64 | 8.00 |
| 9 | PATHWAYS IN CANCER | 0.1196 | 0.4269 | 2.60 | 5.52 |
| 10 | HEDGEHOG SIGNALING PATHWAY | 0.1211 | 0.4269 | 8.04 | 16.97 |
| 11 | ACUTE MYELOID LEUKEMIA | 0.1432 | 0.4269 | 6.70 | 13.02 |
| 12 | NON-SMALL CELL LUNG CANCER | 0.1511 | 0.4269 | 6.31 | 11.93 |
| 13 | CYTOSOLIC DNA-SENSING PATHWAY | 0.1724 | 0.4269 | 5.45 | 9.59 |
| 14 | ERBB SIGNALING PATHWAY | 0.1743 | 0.4269 | 5.39 | 9.41 |
| 15 | COMPLEMENT AND COAGULATION CASCADES | 0.1819 | 0.4269 | 5.13 | 8.75 |
| 16 | MORPHINE ADDICTION | 0.1857 | 0.4269 | 5.02 | 8.45 |
| 17 | PI3K-AKT SIGNALING PATHWAY | 0.1965 | 0.4269 | 2.52 | 4.11 |
| 18 | AGE-RAGE SIGNALING PATHWAY IN DIABETIC COMPLICATIONS | 0.2061 | 0.4269 | 4.46 | 7.04 |
| 19 | AMOEBIASIS | 0.2061 | 0.4269 | 4.46 | 7.04 |
| 20 | PROTEIN DIGESTION AND ABSORPTION | 0.2098 | 0.4269 | 4.37 | 6.82 |
| 21 | MRNA SURVEILLANCE PATHWAY | 0.2098 | 0.4269 | 4.37 | 6.82 |
| 22 | SYSTEMIC LUPUS ERYTHEMATOSUS | 0.2134 | 0.4269 | 4.28 | 6.61 |
| 23 | GLUTAMATERGIC SYNAPSE | 0.2279 | 0.4359 | 3.97 | 5.88 |
| 24 | RELAXIN SIGNALING PATHWAY | 0.2525 | 0.4629 | 3.53 | 4.85 |

| Index | Name | P-value | Adjusted p-value | Odds Ratio | Combined score |
|---|---|---|---|---|---|
| 25 | OOCYTE MEIOSIS | 0.2696 | 0.4667 | 3.26 | 4.28 |

**Extended Data Table 3. Expression of select genes in fetal intestine**

| Common name | E14.5 Rep1 | E14.5 rep2 | E15.5 rep1 | E15.5 rep2 | E16.5 rep1 | E16.5 rep2 | PND0 rep1 | PND0 rep2 |
|---|---|---|---|---|---|---|---|---|
| Adgrg6 | 1.62 | 1.81 | 1.47 | 1.4 | 0.82 | 0.88 | 0.66 | 0.68 |
| Bex4 | 100.81 | 103.84 | 133.2 | 151.04 | 144.94 | 151.34 | 152.17 | 187.16 |
| Bhlhb9 | 24.11 | 22.29 | 18.84 | 20.49 | 14.8 | 12.84 | 6.5 | 5.86 |
| Cadps | 1.22 | 1.26 | 0.81 | 0.93 | 0.52 | 0.43 | 0.43 | 0.45 |
| Chd3 | 32.8 | 27.96 | 25.46 | 22.79 | 21.18 | 19.54 | 12.46 | 15.49 |
| Col4a1 | 97.28 | 100.1 | 79.59 | 84.37 | 63.8 | 81.11 | 52.49 | 52.67 |
| Dnajc6 | 1.11 | 0.81 | 1 | 1.1 | 0.68 | 0.75 | 0.78 | 0.59 |
| Eid2 | 1.77 | 1.7 | 2.61 | 4.15 | 0.46 | 0.58 | 2.01 | 2.38 |
| Erbb4 | 0.03 | 0.04 | 0.03 | 0.01 | 0 | 0.04 | 0 | 0.01 |
| F5 | 0.32 | 0.31 | 0.25 | 0.24 | 0.19 | 0.21 | 0.15 | 0.26 |
| Fam20c | 6.31 | 5.48 | 4.56 | 4.05 | 1.97 | 2.06 | 1.61 | 2.17 |
| Fndc1 | 2.98 | 4.92 | 6.55 | 8.11 | 7.91 | 8.75 | 10.09 | 11.87 |
| Fras1 | 7.58 | 8.02 | 3.46 | 3.24 | 2.36 | 2.49 | 0.95 | 1.03 |
| Gata5 | 4.2 | 4.37 | 4.95 | 5.08 | 6.15 | 6.8 | 13.56 | 15.17 |
| Gata5os | 0.21 | 0.12 | 0.44 | 0.22 | 0.48 | 0.62 | 1.34 | 2.44 |
| Gpc2 | 2.65 | 1.84 | 8.72 | 7.82 | 7.8 | 6.57 | 0.96 | 0.94 |
| Grk3 | 4.98 | 4.1 | 3.56 | 3.45 | 2.87 | 3.2 | 1.47 | 1.77 |
| Gsg1l | 0.66 | 0.55 | 0.33 | 0.39 | 0.29 | 0.4 | 0.33 | 0.23 |
| Gspt2 | 3.44 | 3.71 | 2.03 | 2.65 | 2.35 | 1.84 | 0.93 | 0.97 |
| Kcnb2 | 2.26 | 2.02 | 1.18 | 0.94 | 0.84 | 0.93 | 0.51 | 0.72 |
| Ltbp2 | 0.35 | 0.58 | 0.94 | 0.43 | 0.49 | 1.09 | 0.56 | 0.65 |
| Mab21l1 | 11.3 | 8.24 | 5.58 | 6.82 | 5.12 | 4.71 | 2.33 | 2.31 |
| Macroh2a2 | 63.51 | 55.94 | 52.28 | 62.3 | 45.95 | 39.28 | 14.93 | 13.53 |
| Mn1 | 6.22 | 5.89 | 4.63 | 5.02 | 4.44 | 4.24 | 1.85 | 1.98 |
| Nebl | 0.35 | 0.53 | 0.63 | 0.25 | 0.38 | 0.28 | 0.14 | 0.2 |
| Onecut2 | 1.66 | 3.01 | 1.84 | 1.86 | 1.83 | 1.41 | 3.47 | 4.26 |
| Peg3 | 77.08 | 84.71 | 58.76 | 63.53 | 34.52 | 36.77 | 9.67 | 10.55 |
| Plxdc2 | 12.29 | 10.65 | 9.93 | 9.98 | 7.39 | 7.83 | 2.75 | 2.9 |
| Ppic | 137.63 | 127.18 | 115.84 | 130.92 | 116.71 | 105.21 | 69.9 | 69.17 |
| Prdm5 | 10.19 | 8.2 | 6.35 | 6.81 | 5.31 | 5.13 | 1.74 | 2.11 |
| Rarb | 14.58 | 12.65 | 6.61 | 7.56 | 4.91 | 4.51 | 1.96 | 1.95 |
| Rbp1 | 58.07 | 56.77 | 36.35 | 38.25 | 30.81 | 25.2 | 16.51 | 16.88 |
| Robo1 | 12.98 | 10.27 | 7.18 | 8.11 | 4.99 | 5.01 | 1.42 | 1.49 |
| Runx1t1 | 11.73 | 11 | 7.25 | 7.23 | 5.28 | 4.92 | 1.81 | 2.03 |
| Sall2 | 8.57 | 8.22 | 5.34 | 5.87 | 4.35 | 4.44 | 1.57 | 1.72 |
| St8sia6 | 0.35 | 0.37 | 0.19 | 0.21 | 0.19 | 0.28 | 0.41 | 0.37 |
| Stag3 | 1.61 | 2.58 | 1.66 | 1.14 | 1.9 | 1.18 | 4.1 | 4.8 |
| Stk26 | 3.87 | 3.31 | 3.51 | 3.81 | 3.1 | 3.24 | 3.94 | 3.69 |
| Sult4a1 | 2.79 | 2.35 | 3.3 | 2.67 | 2.63 | 2.56 | 2.59 | 2.26 |
| Tacstd2 | 5 | 3.55 | 3.35 | 4.62 | 3.7 | 2.66 | 0.11 | 0.07 |

| | | | | | | | | |
|---|---|---|---|---|---|---|---|---|
| Tiam1 | 12.52 | 11.47 | 7.79 | 7.22 | 4.68 | 4.91 | 2.09 | 2 |
| Trim35 | 63.06 | 62.16 | 53.56 | 56.18 | 47.65 | 46.88 | 21.51 | 20.99 |
| Zcchc3 | 20.76 | 20.65 | 19.92 | 23.42 | 12.38 | 12.43 | 10.18 | 11.5 |
| Zfp105 | 10.45 | 9.45 | 7.03 | 7.88 | 6 | 5.21 | 2.28 | 3.04 |
| Zfp334 | 5.52 | 4.97 | 2.86 | 2.86 | 2.16 | 2.1 | 0.58 | 0.52 |
| Zfp518b | 9.93 | 8.81 | 7.72 | 8.16 | 5.93 | 5.62 | 1.57 | 1.53 |

Expression value of genes identified in the text were queried in a publicly available dataset of fetal intestinal RNA expression[1]. Data are provided as FPKM as reported in the public archive. Value less than 0.5 FPKM are highlighted in red.

1. He, P. *et al*. The changing mouse embryo transcriptome at whole tissue and single-cell resolution. *Nature* **583**, 760-767 (2020).

Extended Data Table 4. Hallmark Pathway and Reactome Enrichment

Top 25 Scoring Enriched Pathways in MSigDB Hallmark 2020

| Index | Name | P-value | Adjusted p-value | Odds Ratio | Combined score |
|---|---|---|---|---|---|
| 1 | Interferon Alpha Response | 7.307e-33 | 3.288e-31 | 17.80 | 1316.99 |
| 2 | Interferon Gamma Response | 2.303e-30 | 5.181e-29 | 8.86 | 604.32 |
| 3 | TNF-alpha Signaling via NF-kB | 2.222e-10 | 3.334e-9 | 4.16 | 92.45 |
| 4 | KRAS Signaling Up | 1.932e-7 | 0.000002174 | 3.41 | 52.67 |
| 5 | p53 Pathway | 6.645e-7 | 0.000005981 | 3.26 | 46.40 |
| 6 | IL-6/JAK/STAT3 Signaling | 0.00004623 | 0.0003399 | 3.97 | 39.66 |
| 7 | Inflammatory Response | 0.00006042 | 0.0003399 | 2.70 | 26.26 |
| 8 | Xenobiotic Metabolism | 0.00006042 | 0.0003399 | 2.70 | 26.26 |
| 9 | Epithelial Mesenchymal Transition | 0.0001665 | 0.0008325 | 2.57 | 22.35 |
| 10 | Allograft Rejection | 0.0004371 | 0.001967 | 2.44 | 18.84 |
| 11 | Coagulation | 0.001949 | 0.007973 | 2.52 | 15.74 |
| 12 | Hypoxia | 0.002590 | 0.009713 | 2.17 | 12.95 |
| 13 | Reactive Oxygen Species Pathway | 0.007212 | 0.02497 | 3.43 | 16.93 |
| 14 | Myogenesis | 0.01240 | 0.03987 | 1.92 | 8.43 |
| 15 | IL-2/STAT5 Signaling | 0.02394 | 0.07183 | 1.80 | 6.73 |
| 16 | Apoptosis | 0.03798 | 0.1068 | 1.81 | 5.92 |
| 17 | Apical Junction | 0.04731 | 0.1183 | 1.67 | 5.10 |
| 18 | Complement | 0.04731 | 0.1183 | 1.67 | 5.10 |
| 19 | Apical Surface | 0.05250 | 0.1244 | 2.63 | 7.76 |
| 20 | Cholesterol Homeostasis | 0.05627 | 0.1266 | 2.15 | 6.18 |
| 21 | Fatty Acid Metabolism | 0.06555 | 0.1405 | 1.69 | 4.61 |
| 22 | Estrogen Response Late | 0.08446 | 0.1658 | 1.55 | 3.83 |
| 23 | Angiogenesis | 0.08473 | 0.1658 | 2.57 | 6.34 |
| 24 | UV Response Dn | 0.1348 | 0.2361 | 1.53 | 3.07 |
| 25 | Estrogen Response Early | 0.1417 | 0.2361 | 1.43 | 2.79 |

Top 25 Scoring Enriched Pathways in Reactome Pathways 2024

| Index | Name | P-value | Adjusted p-value | Odds Ratio | Combined score |
|---|---|---|---|---|---|
| 1 | Interferon Alpha Beta Signaling | 4.582e-12 | 4.476e-9 | 8.22 | 214.54 |
| 2 | Interferon Signaling | 9.277e-10 | 4.532e-7 | 3.42 | 71.04 |
| 3 | Cytokine Signaling in Immune System | 1.547e-9 | 5.038e-7 | 2.30 | 46.57 |
| 4 | NFE2L2 Regulating Anti-Oxidant Detoxification Enzymes | 5.983e-8 | 0.00001461 | 18.60 | 309.42 |
| 5 | Interferon Gamma Signaling | 1.285e-7 | 0.00002510 | 4.95 | 78.47 |
| 6 | Immune System | 0.00002232 | 0.003634 | 1.50 | 16.09 |
| 7 | Nuclear Events Mediated by NFE2L2 | 0.00004623 | 0.006452 | 3.97 | 39.66 |
| 8 | Type I Hemidesmosome Assembly | 0.00007898 | 0.009645 | 17.16 | 162.06 |
| 9 | Glycerophospholipid Catabolism | 0.0001458 | 0.01583 | 27.42 | 242.23 |
| 10 | Glutathione Conjugation | 0.0002476 | 0.02419 | 5.69 | 47.25 |
| 11 | Laminin Interactions | 0.0003677 | 0.02923 | 6.27 | 49.61 |
| 12 | Common Pathway of Fibrin Clot Formation | 0.0003929 | 0.02923 | 7.72 | 60.57 |
| 13 | Cell Junction Organization | 0.0004189 | 0.02923 | 2.99 | 23.23 |
| 14 | KEAP1-NFE2L2 Pathway | 0.0004189 | 0.02923 | 2.99 | 23.23 |
| 15 | Heme Degradation | 0.0006144 | 0.04002 | 9.35 | 69.18 |
| 16 | Extracellular Matrix Organization | 0.0008533 | 0.05055 | 2.06 | 14.53 |
| 17 | Platelet Degranulation | 0.0009082 | 0.05055 | 2.75 | 19.24 |
| 18 | Collagen Formation | 0.0009313 | 0.05055 | 3.18 | 22.19 |
| 19 | Interleukin-10 Signaling | 0.001156 | 0.05943 | 4.34 | 29.36 |
| 20 | Response to Elevated Platelet Cytosolic Ca2+ | 0.001346 | 0.06576 | 2.63 | 17.39 |
| 21 | Biosynthesis of Specialized Proresolving Mediators (SPMs) | 0.001456 | 0.06774 | 7.35 | 48.01 |
| 22 | Non-integrin membrane-ECM Interactions | 0.001528 | 0.06784 | 3.71 | 24.08 |
| 23 | Dissolution of Fibrin Clot | 0.002380 | 0.1011 | 9.14 | 55.20 |
| 24 | Nicotinate Metabolism | 0.002712 | 0.1104 | 4.94 | 29.20 |
| 25 | Biological Oxidations | 0.003744 | 0.1463 | 2.05 | 11.46 |
